# Temporal integration is a robust feature of perceptual decisions

**DOI:** 10.1101/2022.10.25.513647

**Authors:** Alexandre Hyafil, Jaime de la Rocha, Cristina Pericas, Leor N. Katz, Alexander C. Huk, Jonathan W. Pillow

## Abstract

Making informed decisions in noisy environments requires integrating sensory information over time. However, recent work has suggested that it may be difficult to determine whether an animal’s decision-making strategy relies on evidence integration or not. In particular, strategies based on extrema-detection or random snapshots of the evidence stream may be difficult or even impossible to distinguish from classic evidence integration. Moreover, such non-integration strategies might be surprisingly common in experiments that aimed to study decisions based on integration. To determine whether temporal integration is central to perceptual decision making, we developed a new model-based approach for comparing temporal integration against alternative “non-integration” strategies for tasks in which the sensory signal is composed of discrete stimulus samples. We applied these methods to behavioral data from monkeys, rats, and humans performing a variety of sensory decision-making tasks. In all species and tasks, we found converging evidence in favor of temporal integration. First, in all observers across studies, the integration model better accounted for standard behavioral statistics such as psychometric curves and psychophysical kernels. Second, we found that sensory samples with large evidence do not contribute disproportionately to subject choices, as predicted by an extrema-detection strategy. Finally, we provide a direct confirmation of temporal integration by showing that the sum of both early and late evidence contributed to observer decisions. Overall, our results provide experimental evidence suggesting that temporal integration is an ubiquitous feature in mammalian perceptual decision-making. Our study also highlights the benefits of using experimental paradigms where the temporal stream of sensory evidence is controlled explicitly by the experimenter, and known precisely by the analyst, to characterize the temporal properties of the decision process.

## INTRODUCTION

Perceptual decision-making is thought to rely on the temporal integration of noisy sensory information on a timescale of hundreds of milliseconds to seconds. Temporal integration corresponds to summing over time the evidence provided by each new sensory stimulus, and optimizes perceptual judgments in face of noise (Bogacz et al. 2006; Gold and Shadlen 2007). A perceptual decision can then be made on the basis of this accumulated evidence, either as some threshold on accumulated evidence is reached, or if some internal or external cue signals the need to initiate a response.

Although many behavioral and neural results are consistent with this integration framework, temporal integration is a feature that has often been taken for granted rather than explicitly tested. Recently, the claim that standard perceptual decision-making tasks rely on (or even frequently elicit) temporal integration has been challenged by theoretical results showing that non-integration strategies can produce behavior that carries superficial signatures of temporal integration (Stine et al. 2020). These signatures include the relationship between stimulus difficulty, stimulus duration and behavioral accuracy, the precise temporal weighting of sensory information on the decisions, and the patterns of reaction times.

Here, we propose new analytical tools for directly assessing integration and non-integration strategies from fixed-duration or variable-duration paradigms where, critically, the experimenter controls the fluctuations in perceptual evidence over time within each trial (discrete-sample stimulus, or DSS). By leveraging these controlled fluctuations, our methods allow us to make direct comparisons between integration and non-integration strategies. We apply these tools to assess temporal integration in data from monkeys, humans and rats that performed a variety of perceptual decision-making tasks with DSS. Applying these analyses to these behavioral datasets yields strong evidence that perceptual decision-making tasks in all three species rely on temporal integration. Temporal integration, a critical element of many major theories of perception at both the neural and behavioral levels, is indeed a robust and pervasive aspect of mammalian behavior. Our results also illuminate the power of targeted stimulus design and statistical analysis to test specific features of behavior.

## RESULTS

### Integration and non-integration models

In a typical perceptual evidence-integration experiment (Figure 1A), an observer is presented in each trial with a time-varying stimulus and must report which of two possible stimulus categories it belongs to. Typical examples include judging whether a dynamic visual stimulus is moving leftwards or rightwards (Yates et al. 2017; Katz et al. 2015); whether the orientation of a set of gratings is more aligned with cardinal or diagonal directions (Wyart et al. 2012); whether a combination of tones is dominated by high or low frequencies (Morillon, Schroeder, and Wyart 2014; Hermoso-Mendizabal et al. 2020; Znamenskiy and Zador 2013); or which of two acoustic streams is more intense or dense (Brunton, Botvinick, and Brody 2013; Pardo-Vazquez et al. 2019). Such paradigms have been used extensively in humans, nonhuman primates and rodents. Here we focus on experiments in which observers report their choice at the end of a period whose duration is controlled by the experimenter (Kiani and Shadlen 2009; Wyart et al. 2012; Brunton, Botvinick, and Brody 2013; Raposo et al. 2012), in contrast to so-called *“*reaction time*”* tasks, in which the observer can respond after viewing as brief a portion of the stimulus as they wish (Roitman and Shadlen 2002; Znamenskiy and Zador 2013; Pardo-Vazquez et al. 2019; Hermoso-Mendizabal et al. 2020).

**Figure 1.**
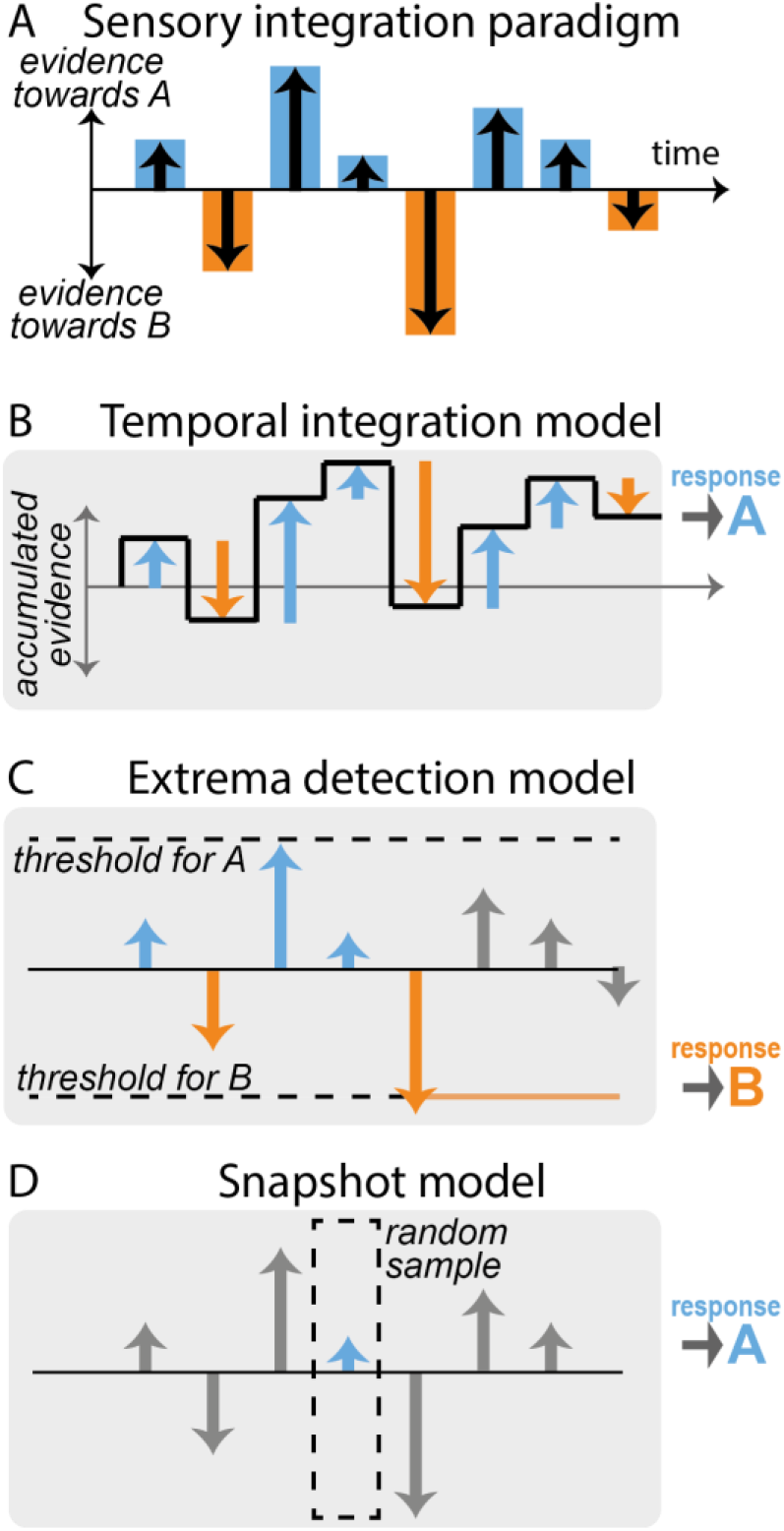
**A**. Schematic of a typical fixed-duration perceptual task with discrete-sample stimuli (DSS). A stimulus is composed of a discrete sequence of n samples (here. *n=6*). The subjects must report at the end of the sequence whether one specific quality of the stimulus was *“*overall*”* leaning more towards one of two possible categones A or B. Evidence in favor of category A or B varies across samples (blue and orange bars). **B**. Temporal integration model. The relative evidence in favor of each category is accumulated sequentially as each new sample is presented (black line), resulting in temporal integration of the sequence evidence. The choice is determined by the end point of the accumulation process: here, the overall evidence in favor of category A is positive, so response A is selected. **C**. Extrema detection model. A decision is made whenever the instantaneous evidence for a given sample (blue and orange arrows) reaches a certain fixed threshold (dotted lines). The selected choice corresponds to the sign of the evidence of the sample that reaches the threshold (here, response B). Subsequent samples are ignored (gray bars). **D**. Snapshot model Here, only one sample is attended. Which sample is attended is determined in each trial by a stochastic policy. The response of the model simply depends on the evidence of the attended sample. Other samples are ignored (gray bars). Variants of the model include attending *K*>*1* sequential samples.

Moreover, we focus on experimental paradigms in which the sensory evidence in favor of each category arrives in a sequence of discrete *samples*. Samples can correspond to motion pulses (Yates et al. 2017), individual gratings (Wyart et al. 2012), acoustic tones (Morillon, Schroeder, and Wyart 2014; Hermoso-Mendizabal et al. 2020; Znamenskiy and Zador 2013) numbers (Bronfman et al. 2015) or symbols representing category probabilities (Yang and Shadlen 2007). We refer to this configuration as the discrete-sample stimulus (DSS) paradigm. In this paradigm, the perceptual evidence provided by each sample can be controlled independently, allowing for detailed analyses of how different samples contribute to the behavioral response. The DSS framework can be contrasted with experiments in which the experimenter specifies only the mean stimulus strength on each trial, and variations in sensory evidence over time are not finely controlled or are not easily determined from the raw spatio-temporal stimulus.

Tasks using the DSS paradigm are classically thought to rely on sequential accumulation of the stimulus evidence (Bogacz et al. 2006), which we refer to here as temporal integration. Figure 1A shows an example stimulus sequence composed of *n* samples that provide differing amounts of evidence in favor of one alternative vs. another (*“*A*”* vs. *“*B*”*). The accumulated evidence fluctuates as new samples are integrated and finishes at a positive value indicating overall evidence for stimulus category A (Figure 1B). This integration process can be formalized by defining the the decision variable or accumulated evidence *x*_*i*_ and its updating dynamics across stimulus samples: *x*_*i*_ = *x*_*i*−1_ + *m*_*i*_ where *m*_*i*_ = *S*_*i*_ + *ε*_*i*_ represents a noisy version of the true stimulus evidence *S*_*i*_ in the *i*-th sample corrupted by sensory noise *ε*_*i*_. The binary decision *r* is simply based on the sign of the accumulated evidence *x*_*n*_ at the end of the sample sequence (composed of *n* samples): *r* = *A* if *x*_*n*_ > 0, and *r* = *B* if *x*_*n*_ < 0. This procedure corresponds to the normative strategy with uniform weighting that maximizes accuracy. For such perfect integration, *x*_*n*_ =*Σ*_*i*_*S*_*i*_ + *Σ*_*i*_*ε*_*i*_, so that the probability of response A is *p*(*r* = *A*) = *Φ*(*Σ*_*i*_*βS*_*i*_) where *Φ* is the cumulative normal distribution function (the normative weight for the stimuli β depends on the noise variance *Var*(*ε*) and the number of samples through 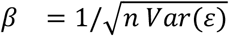. Departures from optimality in the accumulation process such as accumulation leak, categorization dynamics, sensory adaptation or sticky boundaries may however yield unequal weighting of the different samples (Yates et al. 2017; Brunton, Botvinick, and Brody 2013; Prat-Ortega et al. 2021; Bronfman, Brezis, and Usher 2016). To accommodate for these, we allowed the model to take any arbitrary weighting of the samples: *p*(*r* = *A*) = *Φ*(*β*_0_ + *Σ*_*i*_*β*_*i*_*S*_*i*_) (see Methods for details). The mapping from final accumulated evidence to choice was probabilistic, to account for the effects of noise from different sources in the decision-making process (Drugowitsch et al. 2016).

Although it has been commonly assumed that observers use evidence integration strategies to perform these psychophysical tasks, recent work has suggested that observers may employ non-integration strategies instead (Stine et al. 2020). Here we consider two specific alternative models. The first non-integration model corresponds to an *extrema-detection* model (Waskom and Kiani 2018; Stine et al. 2020; Ditterich 2006). In this model, observers do not integrate evidence across samples but instead base their decision on extreme or salient bits of evidence. More specifically, the observer commits to a decision based on the first sample *i* in the stimulus sequence that exceeds one of the two symmetrical thresholds, i.e. such that |*m*_*i*_| ≥ *θ*. In our example stimulus, the first sample that reaches this threshold in evidence space is the fifth sample, which points towards stimulus category B, so response B is selected (Figure 1C). This policy can be viewed as a memory-less decision process with sticky bounds. If the stimulus sequence contains no extreme samples, so that neither threshold is reached, the observer selects a response at random. (Following (Stine et al. 2020), we also explored an alternative mechanism where in such cases the response is based on the last sample in the sequence).

The second non-integration model corresponds to the *snapshot model* (Stine et al. 2020; Pinto et al. 2018). In this model, the observer attends to only one sample *i* within the stimulus sequence, and makes a decision based solely on the evidence from the attended sample: *r* = *A* if *m*_*i*_ > 0, and *r* = *B* if *m*_*i*_ < 0. The position in the sequence of the attended sample is randomly selected on each trial. In our example, the fourth sample is randomly selected, and since it contains evidence towards stimulus category A, response A is selected (Figure 1D). We considered variants of this model that gave it additional flexibility, including: allowing the prior probability over the attended sample to depend on its position in the sequence using a non-parametric probability mass function estimated from the data; allowing for deterministic vs. probabilistic decision-making rule based on the attended evidence; including attentional lapses that were either fixed to 0.02 (split equally between leftward and rightward responses) or estimated from behavioral data. We finally considered a variant of the snapshot where the decision was made based on a sub-sequence of *K* consecutive samples within the main stimulus sequence (1 ≤ *K* < *n*), rather than based on a single sample.

### Standard behavioral statistics favor integration accounts of pulse-based motion perception in primates

To compare the three decision-making models defined above (i.e., temporal integration, extrema-detection, snapshots), we first examined behavioral data from two monkeys performing a fixed-duration motion integration task (Yates et al. 2017). In this experiment, each stimulus was composed of a sequence of 7 motion samples of 150 ms each where the motion strength towards left or right was manipulated independently for each sample. At the end of the stimulus sequence, monkeys reported with a saccade whether the overall sequence contained more motion towards the left or right direction. The animals performed 72137 and 33416 trials for monkey N and monkey P respectively, allowing for in-depth dissection of their response patterns.

We fit the three models (and their variants) to the responses for each animal individually (see Supplementary Figure 1 for estimated parameters for the different models). We then simulated the fitted model and computed, for simulated and experimental data, the psychophysical kernels capturing the weights of the different sensory samples based on their position in the stimulus sequence (Figure 2B). Psychophysical kernels were non-monotonic and differed in shape between the two animals, probably reflecting the complex contributions of various dynamics and sub-optimalities along the sensory and decision pathways (Yates et al. 2017).

**Figure 2.**
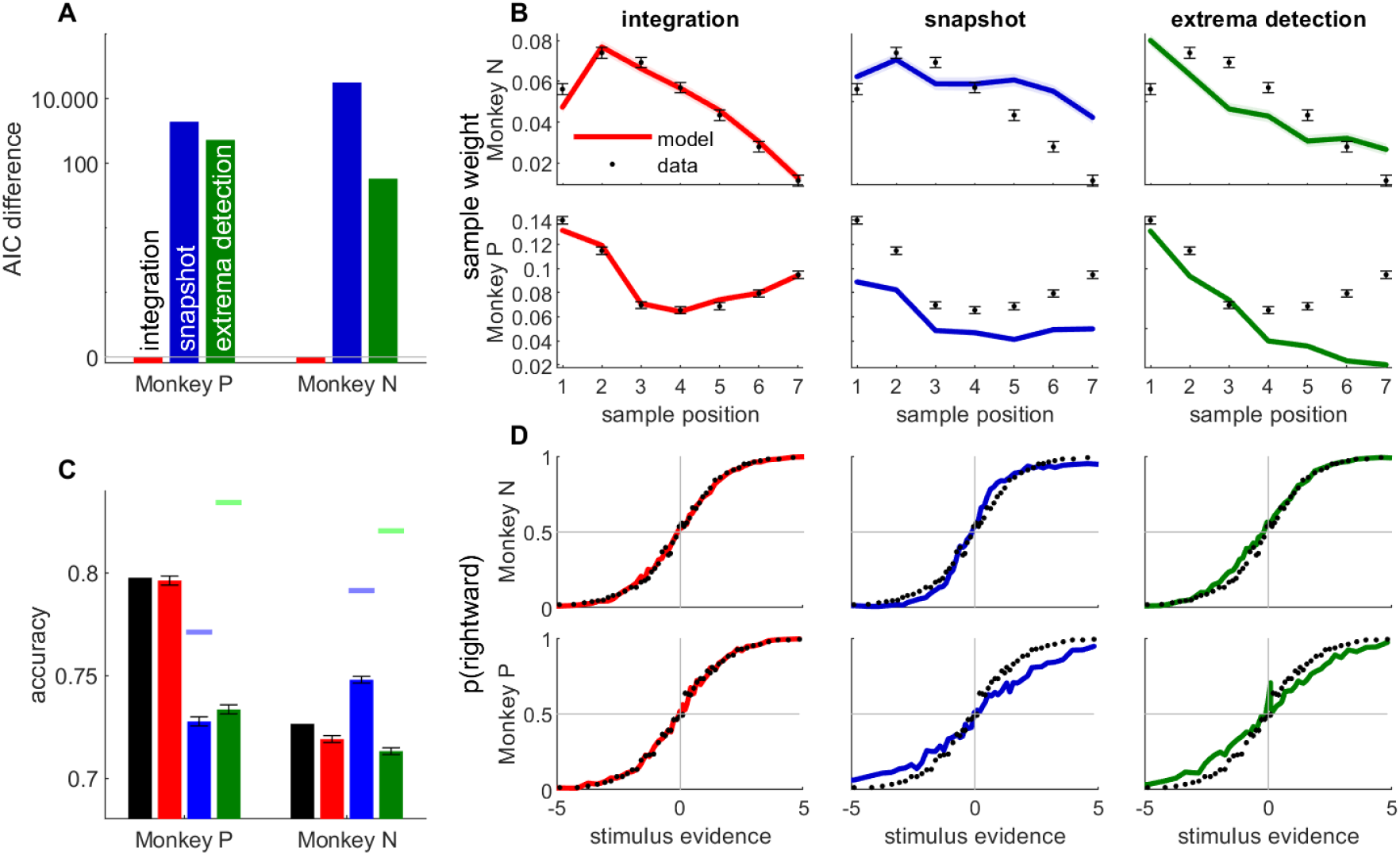
The integration model better described monkey behavior than non-integration models. **A**. Difference between AIC of models (temporal integration: red bar; snapshot model: blue; extrema-detection model: green) and temporal integration model for each monkey. Positive values indicate poorer fit to data. **B**. Psychophysical kernels for behavioral data (black dots) vs. simulated data from temporal integration model (left panel, red curve), snapshot model (middle panel, blue curve) and extrema-detection model (right panel, green curve) for the two animal (monkey N: top panels; monkey P: bottom panels). Each data point represents the weight of the motion pulse at the corresponding position on the animal/model response. Error bars and shadowed areas represent the standard error of the weights for animal and simulated data, respectively. **C**. Accuracy of animal responses (black bars) vs simulated data from fitted models (colour bars), for each monkey. Blue and green marks indicate the maximum performance for the snapshot and extrema-detection models, respectively. Error bars represent standard error of the mean. **D**. Psychometric curves for animal (black dots) and simulated data (colour lines) for monkey N, representing the proportion of rightward choices per quantile of weighted stimulus evidence.

The temporal profile of the kernel was perfectly matched by the integration model, almost by design, as we gave full flexibility to the model to adjust the sample weights. The snapshot model was provided with similar flexibility, as the prior probability of attending each sample could be fully adjusted to the monkey decisions. Surprisingly, however, the snapshot model could not match the experimental psychophysical kernel as accurately. It consistently underestimated the magnitude of weighting in monkey P (Figure 2B, bottom row). The extrema-detection model was not endowed with such flexibility of sensory weighting. On the contrary, since the decision was based on the first sample in the sequence reaching a certain criterion, this inevitably generates a primacy effect in the psychophysical kernels - or at best a flat weighting (Stine et al. 2020). The model thus failed to capture the non-monotonic psychophysical kernels from animal data.

Next, we looked at the psychometric curves and choice accuracy predictions of each fitted model (Figure 2C-D). Stine and colleagues have argued that integration and non-integration models can capture the psychometric curves equally well (Stine et al. 2020). For both animals, the accuracy and psychometric curves were accurately captured by the integration model. In line with Stine and colleagues, we also found that both non-integration models could reproduce the shape of the psychometric curve in monkey N, although the quantitative fit was always better for the integration than non-integration models. By contrast both non-integration models failed to capture the psychometric curve for monkey P (Figure 2B, bottom row). More systematically, the overall accuracy, which is an aggregate measure of the psychometric curve, clearly differs between models, as the accuracy of the non-integration models systematically deviated from animal data for both animals (Figure 2C). In other words, all models produce the same type of psychometric curves up to a scaling factor, and this scaling factor (directly linked to the model accuracy) is key to differentiate model fits. For the snapshot model in monkey P, this discrepancy was explained because the model, limited to using one stimulus sample, could not reach the performance of the model (compare the maximum accuracy of the model indicated by the blue mark with the accuracy of the animal), as the snapshot model is limited to making decisions based on one sensory sample only. (This also explains why the psychophysical kernel of the snapshot model underestimated the true kernel in monkey P). For the extrema-detection model in monkey P and for both non-integration models in the other animal (monkey N) and for the extrema-detection model, the model accuracy is not bounded below the subject*’*s accuracy. In such cases, the model can produce better-than-observed accuracy for certain parameter ranges, but these are not the parameters found by the maximum likelihood procedure, probably because they produce a pattern of errors that is inconsistent with the observed pattern of errors. This indicates an inability of the models to match the pattern of errors of the animal (see Discussion).

Finally, we assessed quantitatively which model provided the best fit, while correcting for model complexity using the Akaike Information Criterion (AIC, Figure 2A). In both monkeys, AIC favored the integration model over the two non-integration models by a very large margin. We also explored whether variants of the extrema-detection and snapshot models could provide a better match to the behavioral metrics considered above (Supp Figure 2 & 3). We found using the AIC metric that the integration model was preferred over all variants of both non-integration models, for both monkeys. Moreover, these model variants could not replicate the psychophysical kernels as well as the integration model did (Supp Figure 2 & 3). In conclusion, while psychometric curves may not always discriminate between integration and non-integration strategies, other metrics including psychophysical kernels, predicted accuracy and quality of fit (AIC) support temporal integration in monkey perceptual decisions. For one model in one monkey (the snapshot model in monkey P), even the simple metric of overall accuracy compellingly supported temporal integration (Fig. 2C). For the other monkey and/or model, where the distinction was less clear, our model-based approach allowed us to leverage these other metrics to reveal strong support for the temporal integration model (Fig. 2A-C). Although these data relies only on two experimental subjects, we show below further evidence supporting the integration model in humans and rats.

### Monkey responses where the largest evidence sample is at odds with the overall stimulus sequence are inconsistent with the extrema-detection model

While formal model comparison leads us to reject the non-integration models in favor of the integration models, it is informative to examine qualitative features of the animal strategies and identify how non-integration models failed to capture them. We started by designing two analyses aimed at testing whether choices were consistent with the extrema-detection model, namely by testing whether choices were strongly correlated with the largest-evidence samples. In the first analysis, we looked at the subset of trials where the evidence provided by the largest-evidence sample in the sequence was at odds with the total evidence in the sequence: we show one example in Figure 3B, where the largest evidence sample points towards response B, while the overall evidence points towards response A. These *‘disagree trials’* represent a substantial minority of the whole dataset: 1865 trials (2.6%) in monkey N, 1831 trials (5.5%) in monkey P. If integration is present, the response of the animal should in general be aligned with the total evidence from the sequence (Figure 3A, red bars). By contrast, if it followed the extrema-detection model (Figure 1C), it should in general follow the largest evidence sample (Figure 3A, green bars). In both monkeys, animal choices were more often than not aligned with the integrated evidence (Figure 3A, black bars), as predicted by the integration model. The responses generated from the extrema-detection model tended to align more with the largest evidence sample, although that behaviour was somehow erratic (for monkey N) due to the large estimated decision noise in the model. This rules out that monkey decisions rely on a memoryless strategy of simply detecting large evidence samples, discarding all information provided by lower evidence samples. Our results complement a previous analysis on disagree trials in this task (Levi et al. 2018), by explicitly comparing monkey behavior to model predictions.

**Figure 3.**
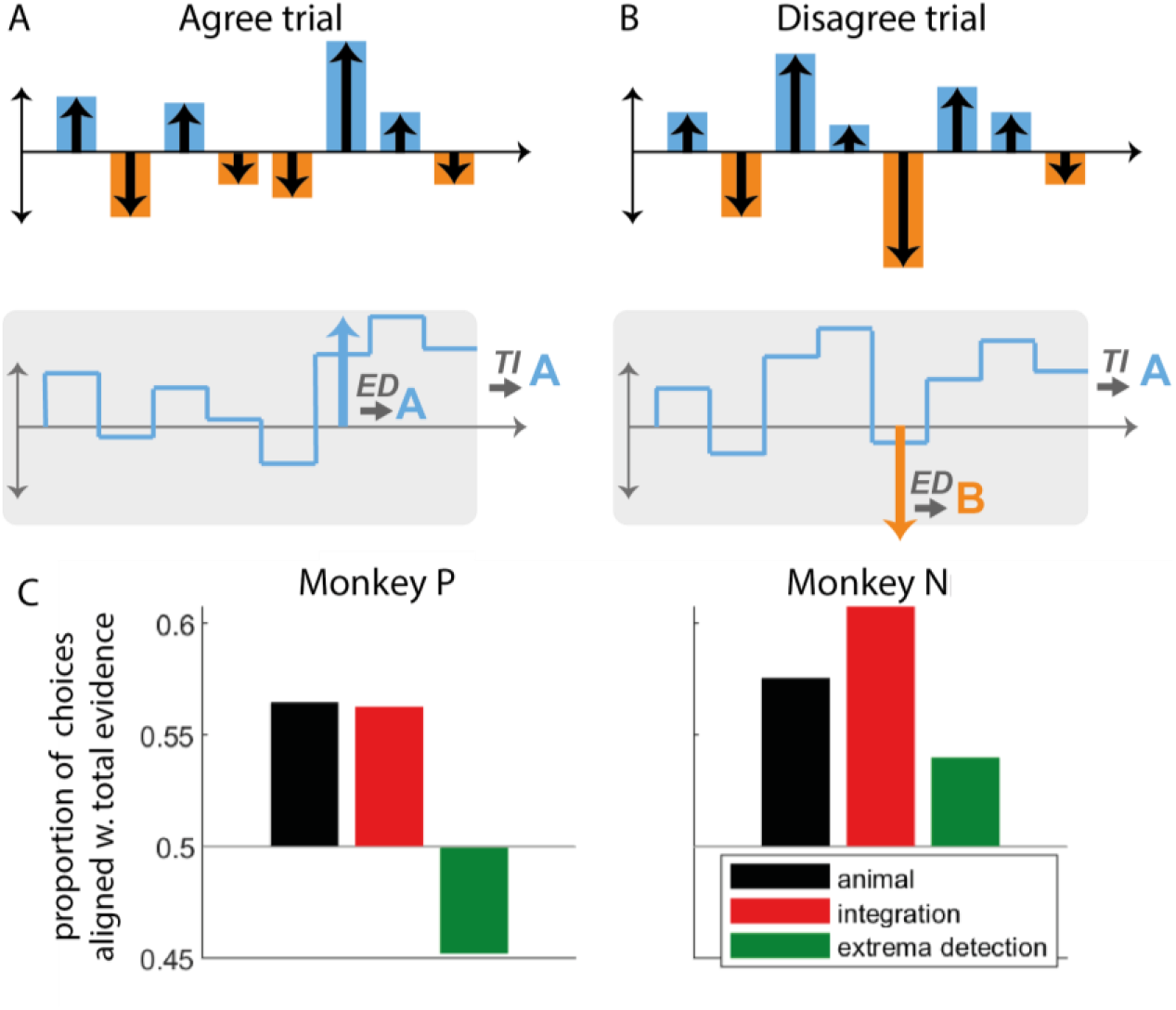
The pattern of animal choices is incompatible with extrema-value based decisions. **A**. Example of an *‘*agree trial*’* where the total stimulus evidence (accumulated over samples) and the evidence from the largest-evidence sample point towards the same response (here, response A). In this case, we expect that temporal integration (TI) and extrema-detection (ED) will produce similar responses (here, A). **B**. Example of a *‘*disagree trial*’*, where the total stimulus evidence and evidence from the largest-evidence sample point towards opposite responses (here A for the former; B for the latter). In this case, we expect that integration and extrema-detection models will produce opposite responses. **C**. Proportion of choices out of all *disagree trials* aligned with total evidence, for animal (black bars), integration (red) and extrema-detection model (green).

We reasoned that the extrema-detection would also leave a clear signature in the *“*subjective weight*”* of the samples, defined as the impact of each sample on the decision as a function of absolute sample evidence (Yang and Shadlen 2007; Waskom and Kiani 2018; Nienborg and Cumming 2007). The extrema-detection model predicts that, in principle, samples whose evidence is below the threshold have little impact on the decision, while samples whose evidence is above the threshold have full impact on the decision. By contrast, the integration model predicts that subjective weight should grow linearly with sample evidence. We estimated subjective weights from monkey choices using a regression method similar in spirit to previous methods (Yang and Shadlen 2007; Waskom and Kiani 2018), taking the form *p*(*r*_*t*_ = *A*) = *σ*(*β*_0_*Σ*_*i*∈[1..*n*]_*β*_*i*_ *f*(*S*_*ti*_)). Here *f* is a function that captures the subjective weight of the sample as a function of its associated evidence. Whereas previous methods estimated subjective weights assuming a uniform psychophysical kernel, our method estimated simultaneously subjective weights *f*(*S*) and the psychophysical kernel *β*, thus removing potential estimation biases due to unequal weighting of sample evidence (see Methods). In both monkeys, we indeed found that the subjective weight depends linearly on sample evidence for low to median values of sample evidence (motion pulse lower than 6), in agreement with the integration model (Supp. Figure 4). Surprisingly however, simulated data of the extrema-detection model displayed the same linear pattern for low to median values of sample evidence. We realized this was due to the very high estimated sensory noise (Supp Fig 1), such that, according to the model, even samples with minimal sample evidence were likely to reach the extrema-detection threshold. In other words, unlike the previous analyses, inferring the subjective weights used by animals was inconclusive as to whether animals deployed the extrema-detection strategy. This somewhat surprising dependency reinforces the importance of validating intuitions by fitting and simulating models (Wilson and Collins 2019).

**Figure 4.**
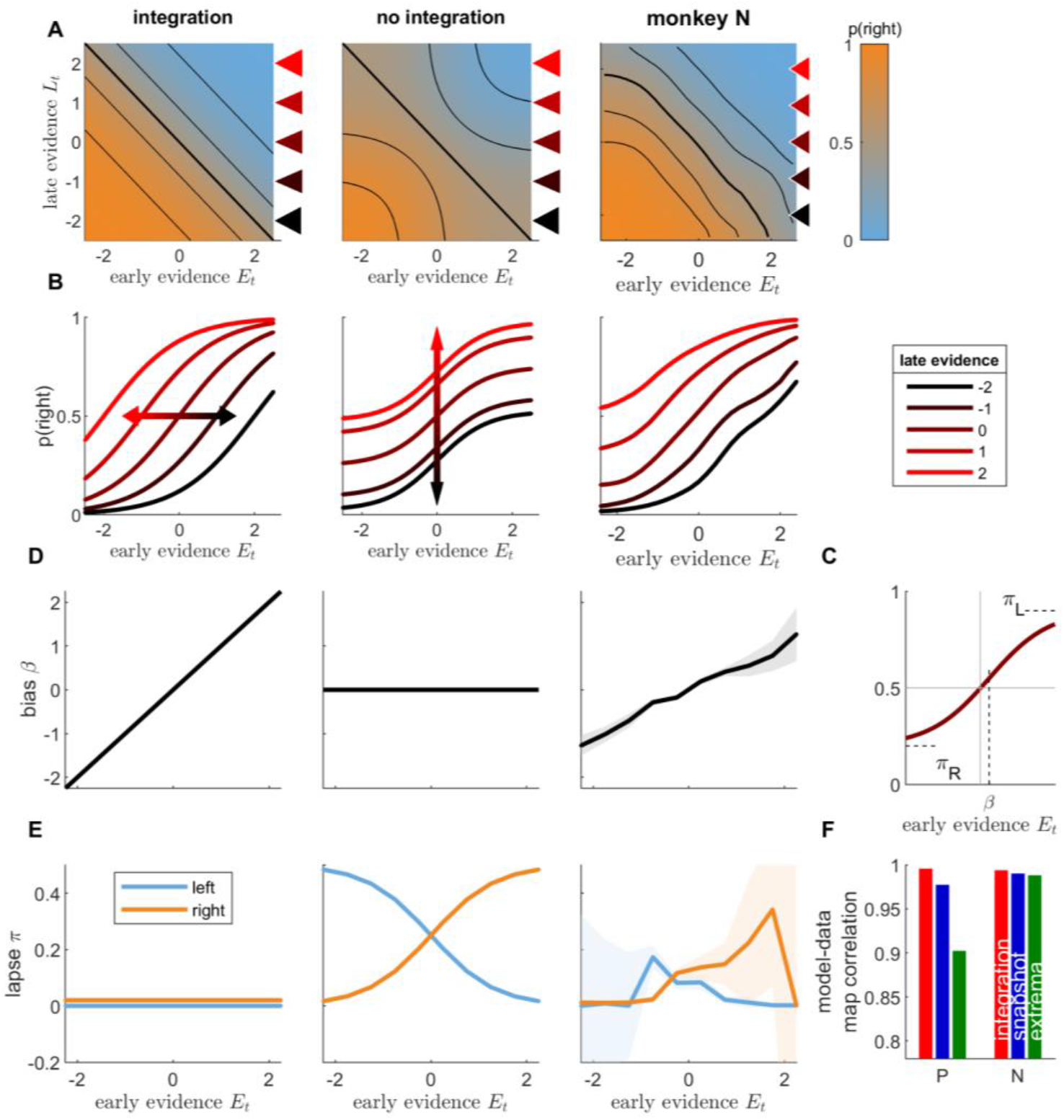
Integration of early and late evidence into animal responses is incompatible with the snapshot model. **A**. Integration map representing the probability of rightward responses (orange: high probability; blue: low probability) as a function of early stimulus evidence *E*_*t*_ and late stimulus evidence *L*_*t*_, illustrated for a toy integration model (where *p*(*right*) = *σ*(*E*_*t*_ + *L*_*t*_); left panel) and a toy non-integration model (*p*(*right*) = 0.5*σ*(*E*_*t*_) + 0.5*σ*(*L*_*t*_); middle panel), and computed for monkey N responses (right panel). Black lines represent the isolines for p(rightwards)=0.15, 0.3, 0.5, 0.7 and 0.85. Conditional psychometric curves representing the probability for rightward response as a function of early evidence *E*_*t*_, for different values of late evidence *L*_*t*_ (see inset for *L*_*t*_values), for toy models and monkey N. The curves correspond to horizontal cuts in the integration maps at *L*_*t*_ values marked by colour triangles in panel A. **C**. Illustration of the fits to conditional psychometric curves. The value of the bias *β*, left lapse *π*_*L*_and right lapse *π*_*R*_are estimated from the conditional psychometric curves for each value of late evidence. **D**. Lateral bias as a function of late evidence for toy models and monkey N. Shaded areas represent standard error of weights for animal data. **E**. Lapse parameters (blue: left lapse; orange: right lapse) as a function of late evidence for toy models and monkey N. **F**. Pearson correlation between integration maps for animal data and integration maps for simulated data, for each animal. Red: integration model; blue: snapshot model; green: extrema-detection model.

### Choice dependence on early and late stimulus evidence show direct evidence for temporal integration

Following model comparisons favoring integration over both snapshot and extrema-detection models, the immediately previous analysis relied on a special subset of trials to provide an additional, and perhaps more intuitive, signature of integration, which ruled out extrema-detection as a possible strategy of either monkey. We next employed another novel analysis specifically designed to tease apart unique signatures of the integration and snapshot models. More specifically, we tested whether decisions were based on the information from only one part of the sequence, as predicted by the snapshot model, or from the full sequence, as predicted by the integration model. To facilitate the analysis, we defined *early evidence E*_*t*_ by grouping evidence from the first three samples in the sequence, and *late evidence L*_*t*_, as the grouped evidence from the last four samples. We then displayed the proportion of rightward responses as a function of both early and late evidence in a graphical representation that we call *integration map* (Figure 4A). A pure integration strategy corresponds to summing early and late evidence equally, which can be formalized as *p*(*r*) = *σ*(*E*_*t*_ + *L*_*t*_), where *σ* is a sigmoidal function. Because this only depends on the sum *E*_*t*_ + *L*_*t*_, the probability of response is invariant to changes in the (*E*_*t*_, *L*_*t*_) space along the diagonal, which leaves the sum unchanged. These diagonals correspond to isolines of the integration map (Figure 4A, left; Supp Figure 5A). In other words, straight diagonal isolines in the integration map reflect the fact that the decision only depends on the sum of evidence *E*_*t*_ + *L*_*t*_. Straight isolines thus constitute a specific signature of evidence integration.

**Figure 5.**
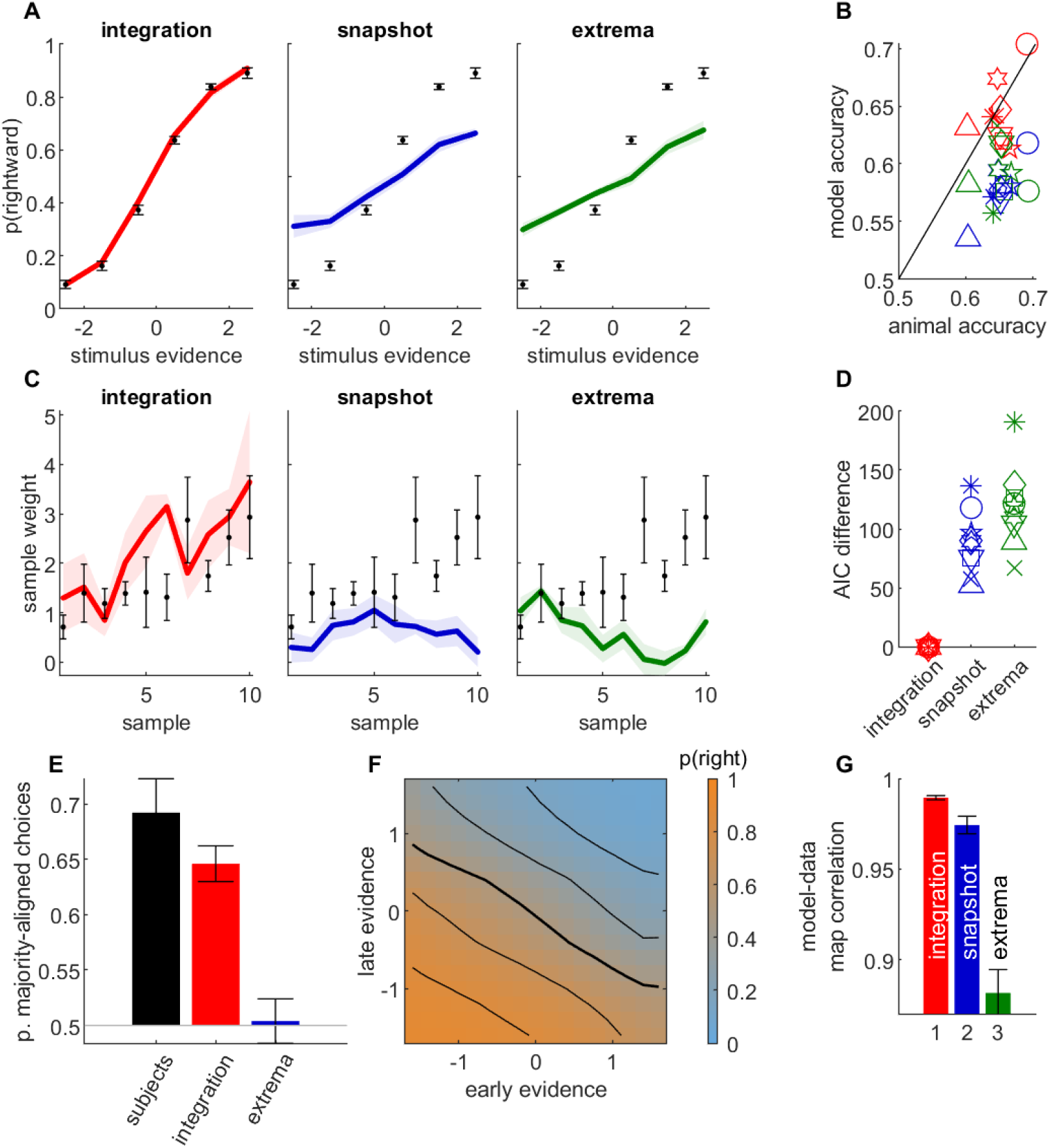
Behavioral data from orientation discrimination task in humans provides further evidence for temporal integration. **A**. Psychometric curves for human data and simulated data, averaged across participants (*n*=9). Legend as in figure 2C. **B**. Simulated model accuracy (y-axis) vs participant accuracy (x-axis) for integration model (red), snapshot model (blue) and extrema-detection model (green). Each symbol corresponds to a participant. **C**. Psychophysical kernel for human data and simulated data, averaged across participants. Legend as in A. **D**. Difference in AIC between each model and the integration model. Legend as in B. **E**. Proportion of choices aligned with total stimulus evidence in disagree trials, for participant data (black bars) and simulated models, averaged over participants. **F**. Integration map for early and late stimulus evidence, computed as in Figure 4A, averaged across participants. **G**. Correlation between integration map of participants and simulated data for integration, snapshot and extrema-detection models, averaged across participants. Colour code as in B. Error bars represent the standard error of the mean across participants in all panels.

We contrasted this integration map with the one obtained from a non-integration strategy (Figure 4A middle panel; Supp Figure 5A). There we assumed that the decision depends either on the early evidence or on the late evidence, as in the snapshot model, with equal probability. This can be formalized as *p*(*r*) = 0.5*σ*(*E*_*t*_) + 0.5*σ*(*L*_*t*_). In this case, if late evidence is null (*σ*(*L*_*t*_) = 0.5) and early evidence is very strong toward the right (*σ*(*E*_*t*_) ≃ 1) the overall probability for rightward response is *p*(*r*) = 0.75. This probability contrasts with that obtained in the integration case where the early evidence would dominate and lead to an overwhelming proportion of rightward responses, i.e. *p*(*r*) ≃ 1. The 25% of leftwards responses yielded by the non-integration model correspond to trials where only the late (uninformative) part of the stimulus is attended and a random response to the left is drawn. More generally, in regions of the space in which either early or late evidence take large absolute values, their corresponding probability of choice saturates to 0 or 1, when that evidence is attended, so the overall response probability becomes only sensitive to the other evidence. As a result, the equiprobable lines bend towards the horizontal and vertical axes (Figure 4A middle). Finally, to compare predictions from both integration and non-integration models to monkey behavior, we plotted the integration maps for both monkeys (Figure 4A, right; Supp Figure 5A). The isolines were almost straight diagonal lines and showed no consistent curvature towards the horizontal and vertical axes. This provides direct evidence that monkey responses depend directly on the sum of early and late evidence— a clear signature of temporal integration.

We derived subsequent tests based on the integration map. We computed conditional psychometric curves as the probability for rightward responses as a function of early evidence *E*_*t*_, conditioned on late evidence value *L*_*t*_ (Figure 4B; Supp Figure 5B). From the integration formula *p*(*r*) = *σ*(*E*_*t*_ + *L*_*t*_), we see that a change in late evidence value corresponds to a horizontal shift of the conditional psychometric curves. By contrast, according to the non-integration formula *p*(*r*) = 0.5*σ*(*E*_*t*_) + 0.5*σ*(*L*_*t*_), conditioning on different values of late evidence adds a fixed value to the response probability irrespective of early evidence, a vertical shift akin to that introduced by lapse responses (Figure 4B middle panel). The conditional psychometric curves for monkeys (Figure 4B right panel; Supp Fig 5 & 6) displayed horizontal shifts as late evidence was changed, consistently with the integration hypothesis. We sought to quantify these shifts in better detail. To this purpose, we fitted each conditional psychometric curve with the formula *p*(*r*) = (1 − *π*_*L*_ − *π*_*R*_) *σ*(*αE*_*t*_ + *β*) + *π*_*R*_, where *π*_*L*_, *π*_*R*_, *α* and *β* correspond to the left lapse, right lapse, sensitivity and lateral bias parameters, respectively (Figure 4C, Supp Fig 5 & 6). The integration model predicts that the bias parameter *β* should vary linearly with *L*_*t*_, while lapse parameters should remain null (Figure 4D, left panel). By contrast, the non-integration model predicts that the horizontal shift parameter *β* should remain constant while left and right lapse parameters (*π*_*L*_, *π*_*R*_) should vary (middle panel), as these lapse parameters correspond to the trials where early evidence is not attended and the response depends simply on late evidence. Both monkeys showed a very strong linear dependence between late evidence and the horizontal shift *β* (Figure 4D, right panel; see also Supp Fig 5), further supporting that late evidence is summed to early evidence.

By contrast, the lapse parameters showed no consistent relationship with late evidence *L*_*t*_ (Figure 4E, right panel). Finally, we directly assessed the similarities between the integration maps from monkey responses and from simulated responses for the three models (integration, snapshot, extrema-detection). The model-data correlation was larger in the integration model than in the non-integration strategies for both monkeys (Figure 4E; unpaired t-test on bootstrapped *r* values: *p<*0.001 for each animal and comparison against extrema-detection and against snapshot model). Overall, integration maps allow to dissect how early and late parts of the stimulus sequence are combined to produce a behavioral response. In both monkeys, these maps carried signatures of temporal integration. For monkey P, the integration model and the data look very similar. For monkey N, there is still a qualitative dependency that deviates from non-integration, but which is not as uniquely matched to the integration strategy (although the imperfect coverage of the two-dimensional space impedes further investigations). Thus, complementing the statistical model tests favoring integration, this richer visualization allows the data to show us that some degree of integration is occurring, albeit not perfect.

### Temporal integration in human visual orientation judgments

Overall, all our analyses converged to support the idea that monkey decisions in a fixed-duration motion discrimination task relied on temporal integration. We explored whether the same results would hold for two other species and perceptual paradigms. We first analyzed the behavioral responses from 9 human subjects performing a variable-duration orientation discrimination task (Cheadle et al. 2014). In each trial, a sequence of 5 to 10 gratings with a certain orientation were shown to the subject, and the subject had to report whether they thought the gratings were overall mostly aligned to the left or to the right diagonal. In this task, the experimenter can control the evidence provided by each sample by adjusting the orientation of the grating. We performed the same analyses on the participant responses than on monkey data. As for monkeys, we found that the integration model nicely captured psychometric curves, participant accuracy and psychophysical kernels (Figure 5A-C, red curves and symbols). By contrast, both non-integration models failed to capture these patterns (Figure 5A-C, blue and green curves and symbols). The accuracy from both models consistently underestimated participant performance: 8 and 6 out of 9 subjects outperformed the maximum performance for the snapshot and extrema-detection models, respectively (Supp. Figure 7). This suggests that human participants achieved such accuracy by integrating sensory evidence over successive samples. Moreover, subjects overall weighted more later samples (Figure 5C), which is inconsistent with the extrema-detection mechanism. A formal model comparison confirmed that in each participant, the integration model provided a far better account of subject responses than either of the non-integration models did (Figure 5D). We then assessed how subjects combined information from weak and strong evidence samples into their decisions, using the same analyses as for monkeys. As predicted by the integration model, but not by the extrema-detection model, humans choices consistently aligned with the total stimulus evidence and not simply with the strongest evidence sample (Figure 5E). Finally, the average integration map for early and late evidence within the stimulus sequence displayed nearly linear diagonal isolines, showing that both were integrated into the response (Figure 5F). Integration maps from participants correlated better with maps predicted by the integration model than with maps predicted by either of the alternative non-integration strategies (Figure 5G; two-tailed t-test on bootstrapped *r* values: p<0.001 for 7 out 9 participants in the integration vs snapshot comparison; in all 9 participants for the integration vs extrema-detection comparison). Overall, these analyses show converging evidence that human decisions in an orientation discrimination task rely on temporal integration.

### Temporal integration in rat acoustic intensity judgments

Finally, we analyzed data from 5 rats performing a fixed-duration auditory task where the animals had to discriminate the side with larger acoustic intensity (Pardo-Vazquez et al. 2019). The relative intensity of the left and right acoustic signals was modulated in sensory samples of 50 ms, so that the stimulus sequence provided time-varying evidence for the rewarded port. The stimulus sequence was composed of either 10 or 20 acoustic samples of 50 ms each, for a total duration of 500 or 1000 ms. We applied the same analysis pipeline as for monkey and human data. The integration model provided a much better account of rat choices than non-integration strategies, based on psychometric curves (Fig. 6A), predicted accuracy (Fig. 6B), psychophysical kernel (Fig. 6C) and model comparison using AIC (Fig. 6D). Similar to humans and monkeys, rats tended to select the side corresponding to the total stimulus evidence and not the largest sample evidence in *“*disagree*”* trials, as predicted by the integration model (Fig. 6E). Finally, the integration map was largely consistent with an integration strategy (Fig. 6F), and correlated more strongly with simulated maps from the integration model (unpaired t-test on bootstrapped *r* values: *p<*0.001 for each animal and comparison against extrema-detection and against snapshot model).

**Figure 6.**
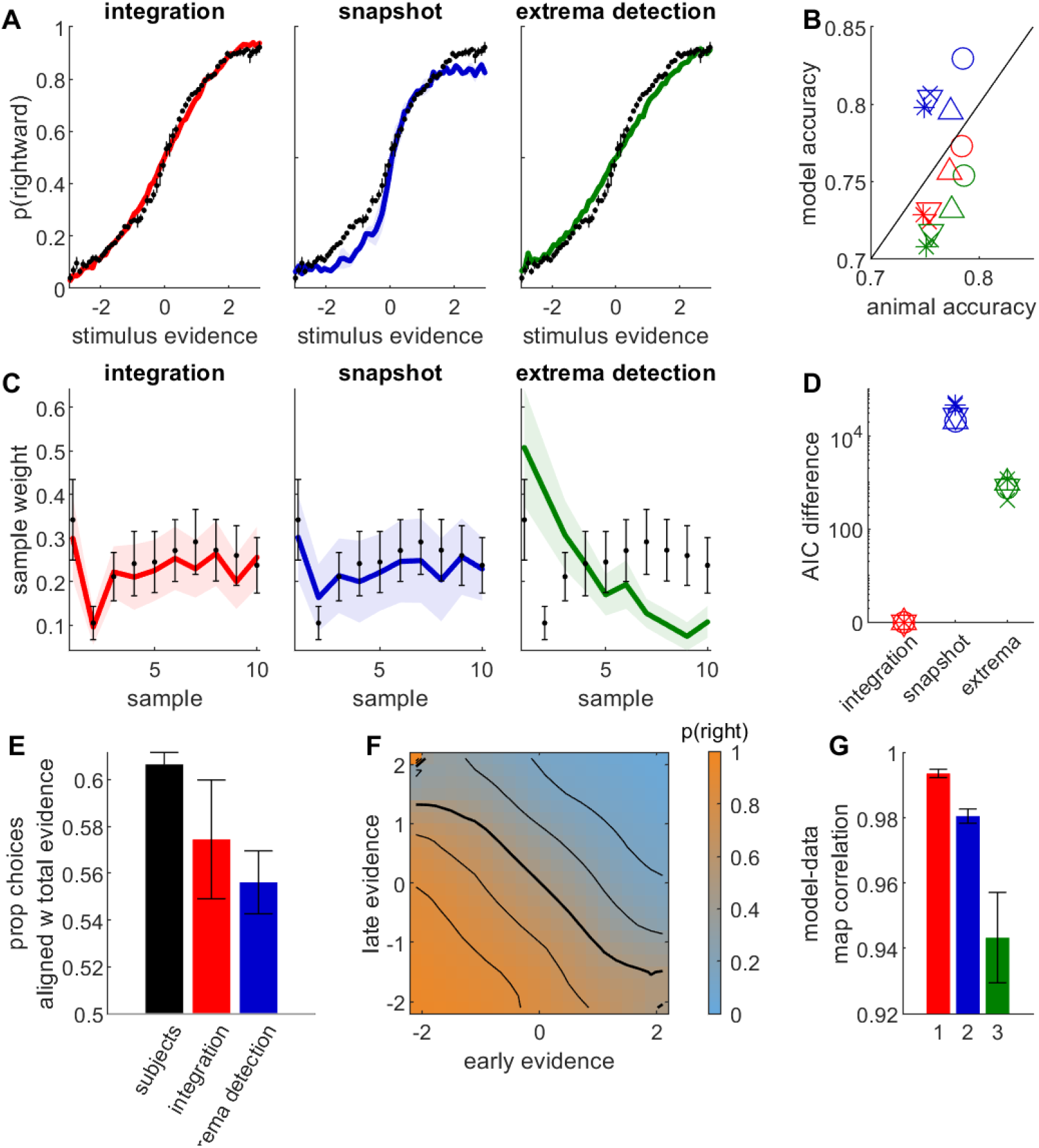
Behavioral data from auditory discrimination task in 5 rats provides further evidence for temporal integration. Rats were rewarded for correctly identifying the auditory sequence of larger intensity (number of samples: 10 or 20; stimulus duration: 500 or 1000 ms). Legend as in Figure 5. Psychophysical kernels are computed only for 10-sample stimuli (in 4 animals). See Supp Figure 8 for psychophysical kernels with 20-sample stimuli.

## DISCUSSION

We investigated the presence of temporal integration in perceptual decisions in monkeys, humans and rats through a series of standard and innovative analyses of response patterns. In all analyses we contrasted predictions from one integration and two non-integration computational models of behavioral responses (Figure 1). For each non-integration model, we considered multiple variants to explore the maximal flexibility offered by each framework to capture animal behavior. For our datasets, evidence in favor of integration was easy to achieve using standard model comparison technique as well as comparing simulated psychometric curves and psychophysical kernels to their experimental counterparts (Figure 2). Our results are in line with previous evidence for temporal integration in perceptual decisions of humans and mice (Pinto et al. 2018; Stine et al. 2020; Waskom and Kiani 2018). Importantly, we also suggest new analyses targeted at revealing specific signatures of temporal integration.

In some cases, we could link the failure of the non-integration model to a fundamental limitation of the model. For example, the extrema-detection model cannot explain the non-monotonic psychophysical kernels of monkeys or the increasing psychophysical kernels in humans. This is because the decision in that mode is based on the first sample to reach a certain fixed criterion, so it will always produce a primacy effect, i.e., a decreasing psychophysical kernel. Although this effect can be small, and in practice yields approximately flat kernels (Stine et al. 2020), it cannot produce increasing or non-monotonic kernels.

Another strong limitation of non-integration models (both the extrema detection and the snapshot model) is that accuracy is limited by the fact that decisions depend on a single sample. We found that that boundary performance (i.e. the maximum performance that a model can reach) was actually lower than subject accuracy for most human participants, *de facto* ruling out these non-integration strategies for these participants. This is consistent to what was observed in a constrast discrimination DSS task where human subjects had to make judgments about image sequences spanning up to tens of seconds each (Waskom and Kiani 2018). It clearly contrasts however with results from (Stine et al. 2020) where the non-integration strategies matched the accuracy of human subjects performing the classical random-dot-motion task. This discrepancy may be related to the different sources of noise in the two paradigms. In DSS tasks, because the sensory evidence provided by the stimulus at each moment is controlled by the experimenter, the unpredictability of human responses essentially stems from internal noise at the level of sensory processing and temporal integration (Waskom and Kiani 2018; Drugowitsch et al. 2016). By contrast, in the random dot motion task (Kiani, Hanks, and Shadlen 2008), which is a non-DSS task because the experimenter does not typically specify differing amounts of motion in each time epoch within a single trial, typically elicits more variable responses due to the presence of stimulus noise. This overall increased noise level leads to a looser relationship between the stimulus condition and the behavioral responses, which can thus be accounted for by a larger spectrum of computational mechanisms. These issues have been addressed by forcing *“*pulses*”* of a certain stimulus strength and/or by performing post hoc analyses to estimate signal and noise (Kiani, Hanks, and Shadlen 2008) but these are partial solutions that DSS paradigms solve by design. This illustrates the benefits of using experimental designs where variability in stimulus information can be fully controlled and parametrized by the experimenter, as these paradigms discriminate more precisely between different models of perceptual decisions.

In at least one monkey, although quantitative metrics such as penalized log-likelihood and fits to psychometric curves clearly pointed to the integration model as the best account to behavior, the qualitative failure modes of the non-integration strategies (especially the snapshot model) was not immediately clear. Although we tried variants for each non-integration model, there remained a possibility that our precise implementation failed to account for monkey behavior but that other possible implementations would. Note that the extrema-detection and snapshot are two of the many possible non-integration strategies. A generic form for non-integration strategies corresponds to a policy that implements position-dependent thresholds on the instantaneous sensory evidence. In this framework, the extrema-dependent model corresponds to the case with a position-independent threshold, while the snapshot model corresponds to a null bound for one sample and infinite bounds for all other samples. To rule out these more complex strategies, we conducted additional analyses that specifically targeted core assumptions of the integration and non-integration strategies.

First, the extrema-detection model fails to account for the data because it predicts that largest-evidence samples should have a disproportionate impact on choices. However, this does not occur, as monkeys and humans tend to respond according to the total evidence and not the single large-evidence sample (Figure 3C and 5E) - see (Levi et al. 2018) for a similar analysis. All non-integration strategies share the property that on each trial the decision should only rely either on the early or the late part of the trial. We thus directly examined the assumptions of integration and non-integration models by assessing how the evidence from the early and late parts of each stimulus sequence is combined to produce a decision. We introduced *integration maps* (Figure 4) to inspect such integration: isolines of the integration maps will be rectilinear if and only if early and late evidence are summed, in other words if and only if temporal integration takes place. Unequal weighting of evidence would still produce rectilinear isolines, albeit with a different angle. By contrast, a non-integration scenario when on each trial only a single piece of evidence contributes to the decision predicts isolines that bend towards the axes. Integration maps from monkey, human and rat subjects nicely matched the predictions of the integration models, proving that their decisions do rely on temporal integration. Note that this innovative analysis technique could be used to probe integration of evidence not only at temporal level but also between different sources of evidence. Indeed, there has been an intense debate about whether sensory information from different spatial locations or different modalities are integrated prior to reaching a decision, or whether decisions are taken separately for each source before being merged, which can be viewed as extensions to the snapshot model (Pannunzi et al. 2015; Otto and Mamassian 2012; Lorteije et al. 2015; Hyafil and Moreno-Bote 2017). Our integration analysis could provide new answers to this old debate.

Integration maps can be computed not only for choice patterns but for any type of behavioral or neural marker of cognition. We computed a neural integration map (Supp. Figure 9) by looking at the average spike activity of Lateral Intra Parietal (LIP) neurons as a function of early and late evidence, for neurons recorded while the monkeys performed the motion discrimination experiment (Yates et al. 2017). The neural integration map clearly showed rectilinear isolines, as predicted by an integration model of neural spiking. By contrast, neural implementations of the snapshot and extrema-detection predicted strongly curved isolines. The activity of LIP neurons correlates with the evidence accumulated over the presentation of the stimulus in favor of either possible choices (Gold and Shadlen 2007). This result shows that the activity of individual LIP neurons indeed reflects the temporal integration of sensory information that drives animal behavior.

We have focused in this study on paradigms where the stimulus duration is fixed by the experimenter, and subjects could only respond after stimulus extinction. Stine et al proposed a method for distinguishing integration from non-integration strategies mixing experiments where stimulus duration is controlled by the experimenter and experiments where the stimulus plays until the subject responds (*“*reaction time paradigms*”*). Our study shows an alternate way to differentiate integration and non-integration strategies that does not require these conditions, and may therefore be applied to existing datasets.

Other studies have shown how integration and non-integration strategies can be disentangled in free reaction-time task paradigms. Specifically, different models make different predictions regarding how the total sample evidence presented before response time should vary with response time (Glickman and Usher 2019; Zuo and Diamond 2019). Glickman and Usher used these predictions to rule out non-integration strategies in a counting task in humans, and Zuo and Diamond found evidence for evidence integration to bound when rats discriminate textures using whisker touches (Zuo and Diamond 2019). Furthermore, decisions in self-paced paradigms are influenced by the sensory evidence from the early part of the stimulus (Winkel et al. 2014), ruling out the proposal that they would only depend on the sensory evidence at the time of decisions (Thura et al. 2012). Of note, the absence of integration seems a more viable strategy when the duration of the stimulus is controlled externally and the benefits of integrating in terms of accuracy might not compensate for its cognitive cost. In free-reaction time paradigms, waiting for a long sequence of samples and selecting its response based on a single sample does not seem a particularly efficient strategy. If the cognitive cost of integration is high, it is more beneficial to interrupt the stimulus sequence early with a rapid response. Such rapid responses are commonly seen and can be attributed either to urgency signals modulating the integration of stimulus evidence (Drugowitsch et al. 2012) or to action initiation mechanisms that time the response after a specific time (e.g. one or two samples) following stimulus onset (Hernández-Navarro et al. 2021). Here, we have shown that even in paradigms where the stimulus duration is controlled by the experimenter, mammals often integrate sensory evidence over the entire stimulus.

In conclusion, we have found strong evidence for temporal integration in perceptual tasks across species (monkeys, humans and rats) and perceptual domain (visual motion, visual orientation and auditory discrimination). Thus, although the time scale of integration can be adapted to the statistics of the environment (Ossmy et al. 2013; Glaze, Kable, and Gold 2015; Kilpatrick et al. 2019), the principle that stimulus evidence is integrated over time appears as a hallmark of perception. This evidence was gathered by leveraging experimentally-controlled sensory evidence at each sensory sample composing a stimulus, and novel model-based statistical analysis. We speculate that temporal integration is a ubiquitous feature of perceptual decisions due to hard-wired neural integrating circuits, such as recurrent stabilizing connectivity in sensory and perceptual areas (Wang 2008; Wimmer et al. 2015).

## METHODS

### Monkey experiment

We present here the most relevant features of the behavioral protocol - see (Yates et al. 2017) for further experimental details. Two adult rhesus macaques (subject N, a 10-year old female; and subject P, a 14-year old male) performed a motion discrimination task. On each trial, a stimulus consisting of a hexagonal grid (5-7 degrees, scaled by eccentricity) of Gabor patches (0.9 cycle per degree; temporal frequency 5 Hz for Monkey P; 7 Hz for Monkey N) was presented. Monkeys were trained to report the net direction of motion in a field of drifting and flickering Gabor elements with an eye movement to one of two targets. Each trial motion stimulus consisted of seven consecutive motion pulses, each lasting 9 or 10 video samples (150 ms or 166 ms; pulse duration did not vary within a session), with no interruptions or gaps between the pulses. The strength and direction of each pulse *S*_*ti*_ for trial *t* and sample *i* was set by a draw from a Gaussian rounded to the nearest integer value. The difficulty of each trial was modulated by manipulating the mean and variance of the Gaussian distribution. Monkeys were rewarded based on the empirical stimulus and not on the stimulus distribution. We analyzed a total of 112 sessions for monkey N and 60 sessions for monkey P, with a total of 72137 and 33416 valid trials, respectively. These sessions correspond to sessions with electrophysiological recordings reported in (Yates et al. 2017) and purely behavioral sessions. All experimental protocols were approved by The University of Texas Institutional Animal Care and Use Committee (AUP-2012-00085, AUP-2015-00068) and in accordance with National Institute of Health standards for care and use of laboratory animals

### Human experiment

9 adult subjects (5 males, 4 females; aged 19-30) performed an orientation discrimination task whereby on each trial they reported in each trial whether a series of gratings were perceived to be mostly tilted clockwise or counterclockwise (Drugowitsch et al. 2016). Each discrete-sample stimulus consisted of five to ten gratings. Each grating was a high-contrast Gabor patch (colour: blue or purple; spatial frequency = 2 cycles per degree; SD of Gaussian envelope = 1 degree) presented within a circular aperture (4 degrees) against a uniform gray background. Each grating was presented during 100 ms, and the interval between gratings was fixed to 300 ms. The angles of the gratings were sampled from a von Mises distribution centered on the reference angle (*α*_0_ = 45 degrees for clockwise sequences, 135 degrees for anticlockwise sequences) and with a concentration coefficient *κ* = 0.3. The normative evidence provided by sample *i* in trial *t* in favor of the clockwise category corresponds to how well the grating orientation *α*_*ti*_ aligns with the reference orientation, i.e. *S*_*ti*_ = 2*κ cos*(2(*α*_*ti*_ −*α*_0_)).

Each sequence was preceded by a rectangle flashed twice during 100 ms (the interval between the flashes and between the second flash and the first grating varied between 300 and 400 ms). The participants indicated their choice with a button press after the onset of a centrally occurring dot that succeeded the rectangle mask and were made with a button press with the right hand. Failure to provide a response within 1000 ms after central dot onset was classified as invalid trial. Auditory feedback was provided 250 ms after participant response (at latest 1100 ms after end of stimulus sequence). It consisted of an ascending tone (400 Hz/800 Hz; 83 ms/167 ms) for correct responses; descending tone (400 Hz/ 400 Hz; 83 ms/167 ms) for incorrect responses; a low tone (400 Hz; 250 ms) for invalid trials.

Trials were separated by a blank interstimulus interval of 1,200-1,600 ms (truncated exponential distribution of mean 1,333 ms). Experiments consisted of 480 trials in 10 blocks of 48. It was preceded with two blocks of initiation with 36 trials each. In the first initiation block, there was only one grating in the sequence, and it was perfectly aligned with one of the reference angles. In the second initiation block, sequences of gratings were introduced, and the difficulty was gradually increased (the distribution concentration linearly decreased from *κ* = 1.2 to *κ* = 0.3). Invalid trials (mean 6.9 per participant, std 9.4) were excluded from all regression analyses. The study was approved by the local ethics committee (approval 2013/5435/I from CEIm-Parc de Salut MAR).

### Rat experiment

Rat experiments were approved by the local ethics committee of the University of Barcelona (Comité d*’*Experimentació Animal, Barcelona, Spain, protocol number Ref 390/14). 5 male Long-Evans rats (no genetic modifications; 350-650g; 8-10 weeks-old at the beginning of the experiment), pair-housed and kept on stable conditions of temperature (23°C) and humidity (60%) with a constant light-dark cycle (12h:12h, experiments were conducted during the light phase). Rats had free access to food, but water was restricted to behavioral sessions. Free water during a limited period was provided on days with no experimental sessions.

Rats performed a fixed-duration auditory discrimination task where they had to classify noisy stimuli based on the intensity difference between the two lateral speakers (Pardo-Vazquez et al. 2019; Hermoso-Mendizabal et al. 2020). A LED on the center port indicated that the rat could start the trial by poking in that center port. After this poke, rats had to hold their snouts in the central port during 300 ms (i.e. fixation). Following this period, an acoustic DSS was played. Rats had to remain in the central port during the entire presentation of the stimulus. At stimulus offset, the center LED went off and rats could then come out of the center port and head towards one of the two lateral ports. Entering the lateral port associated with the speaker that generated the larger sound intensity led to a reward of 24 µl of water (correct responses), while entering the opposite port lead to a 5 s timeout accompanied with a bright light during the entire period (incorrect responses). If rats broke fixation during the pre-stimulus fixation period or during the stimulus presentation, the sound was interrupted, the center LED remained on, and the rat had to initiate a new trial starting by center fixation followed by a new stimulus. Fixation breaks were not included in any of the analyses. Stimulus duration was 0.5 s (10 samples) or 1 s (20 samples). Two rats performed 0.5-second stimuli only (77810 and 54803 valid trials, respectively); one rat performed 1 s stimuli only (42474 valid trials); the remaining two rats performed a mixture of 0.5 and 1 s stimuli trials randomly interleaved (5016 trials and 65212 valid trials, respectively for one animal; 7374 and 38829 trials for the other animal). In each trial *k* one stimulus *S*^*X*^_*k*_(*t*) was played in each speaker (*X*=R for the Right speaker and *X*=L for the Left speaker). Each stimulus was an amplitude modulated (AM) broadband noise defined by 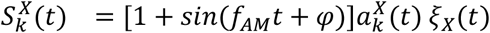 where *f*_AM_=20 Hz (sensory samples lasted 50 ms), the phase delay *φ* = 3*π*/2 and *ξ*_*X*_(*t*) were broadband noise bursts. The amplitudes of each sound in each frame were 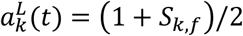 and 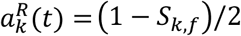 with *S*_k,f_(*t*) being the instantaneous evidence that was drawn independently in each frame *f* from a transformed Beta distribution with support [-1,1]. With this parametrization of the two sounds the sum of the two envelopes was constant in all frames *a*^*L*^_*k*_(*t*)+*a*^*R*^_*k*_(*t*)=1. There were 7 × 5 stimulus conditions, each defined by a Beta distribution, spanning 7 mean values (−1, -0.5, -0.15, 0, 0.15, 0.5 and 1) and 5 different standard deviations (0, 0.11, 0.25, 0.57 and 0.8). In around the first half of the sessions, only sample sequences in which the total stimulus evidence matched the targeted nominal evidence were used. This effectively introduced weak correlations between samples. In the second half of the sessions, this condition was removed and samples in each stimulus were drawn independently from the corresponding Beta distribution.

### Integration model

The integration model for human participants corresponds to a logistic regression model, where the probability of selecting the right choice *p*(*r*_*t*_) at trial *t* depends on the weighted sum of the sample evidence: *p*(*r*_*t*_) = *σ*(*β*_0_ + *Σ*_*i*∈[1..*n*]_*β*_*i*_*S*_*ti*_), where *β*_0_ is a lateral bias, *S*_*ti*_ is the signed sample evidence at sample *i* ;*β*_*i*_ is the sensory weight associated with the *i*th sample in the stimulus sequence; and *σ*(*x*) = (1 + *e*^−*x*^)^−1^is the logistic function. The vector *β*_*i*_*’*s allowed to capture different shapes of psychophysical kernels (e.g. primacy effects, recency effects) which can emerge due to a variety of suboptimalities in the integration process (leak, attractor dynamics, sticky bounds, sensory after-effects, etc.) (Brunton, Botvinick, and Brody 2013; Yates et al. 2017; Prat-Ortega et al. 2021; Bronfman, Brezis, and Usher 2016).

For the monkey and rat data, we included a session-dependent modulation gain *γ*_*t*_ to capture the large variations in performance in monkeys across the course of sessions (see Supp Figure 1A):

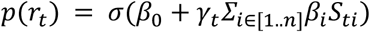

This model corresponds to a bilinear logistic regression model which pertains to the larger family of Generalized Unrestricted Models (GUMs) (Adam and Hyafil 2020). Parameters (*β, γ*) were fitted using the Laplace approximation as described in (Adam and Hyafil 2020). The modulation gain was omitted when applied to human data, yielding a classical logistic regression model.

### Snapshot model

In the snapshot model, decisions are based on each trial based upon a single sample. The model also includes the possibility for left and right lapses. In each trial, the attended sample is drawn from a multinomial distribution of parameters (*π*_1_,.. *π*_*n*_, *π*_*L*_, *π*_*R*_), where the first terms *π*_*i*_ (1 ≤ *i* ≤ *n*) correspond to the probability of attending sample *i*, and *π*_*L*_and *π*_*R*_correspond to the probability of left and right lapses, respectively. Upon selecting sample *i*, the probability for selecting the right choice is given by the function *H*_*i*_(*S*_*t*_). In the deterministic version of the model, *H*_*i*_ is simply determined by the sign of the *i-*th sample evidence:*H*_*i*_(*S*_*t*_) = 1 if *S*_*ti*_ > 0, *H*_*i*_(*S*_*t*_) = 0 if *S*_*ti*_ < 0, and *H*_*i*_(*S*_*t*_) = 0.5 if *S*_*ti*_ = 0 (i.e. random guess if the sample has null evidence). We also define similar functions for lapse responses: *H*_*R*_(*S*) = 1and *H*_*L*_(*S*) = 0, irrespective of the stimulus. In the non-deterministic version of the model, the probability *H*_*i*_(*S*_*ti*_) is determined by a logistic function of the attended sample evidence *H*_*i*_(*S*_*t*_) = *σ*(*β*_*i*_*S*_*ti*_) where *β*_*i*_ describes a sensitivity parameter. The deterministic case can be viewed as the limit of the non-deterministic case when all sensitivity parameters *β*_*i*_ diverge to +∞, i.e. when sensory and decision noise are negligible.

The overall probability for selecting right choice (marginalizing over the attended sample, which is a hidden variable) can be captured by a mixture model :

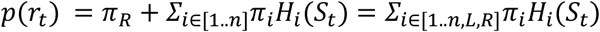

The mixture coefficients *π*_*i*_ (*i* = 1,.. *n, L, R*) are constrained to be non-negative and sum up to In the non-deterministic model, the parameters also include sensitivity parameters *β*_*i*_.The model is fitted using Expectation-Maximization (Bishop 2006). In the Expectation step, we compute the responsibility *z*_*ti*_, i.e. the posterior probability that the sample *i* was attended at trial *t* (for *i=L, R*, the probability that the trial corresponded to a lapse trial):

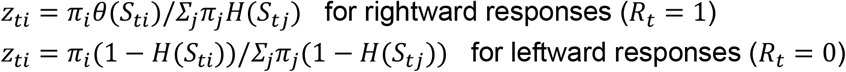

In the Maximization step, we update the value of the parameters by maximizing the Expected Complete Log-Likelihood (ECLL): *Q*(*π, β*) = *Σ*_*ti*_*z*_*ti*_*log p*(*r*_*t*_; *π, β*). Maximizing over the mixture coefficients with the unity-sum constraint provides the classical update: *π*_*i*_ = *Σ*_*ti*_*z*_*ti*_/*N*, where *N* is the total number of trials. In the non-deterministic model, maximizing the ECLL over sensitivity parameters is equivalent to fitting a logistic regression model with weighted coefficients *z*_*ti*_, which is a convex problem. Best fitting parameters can be found using Newton-Raphson updates on the parameters:

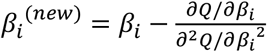

with

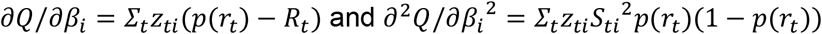

To speed up the computations, in each M step, we only performed one Newton-Raphson update for each sensitivity parameter, rather than iterating the updates fully until convergence. The EM procedure was run until convergence, assessed by an increment in the log-likelihood *L*(*π, β*) of less than 10^−9^ after one EM iteration. The log-likelihood for a given set of parameters is given by *L*(*π, β*) = *Σ*_*t*_*log p*(*r*_*t*_). The EM iterative procedure was repeated with 10 different initializations of the parameters to avoid local minima.

Note that for monkey and rat data, since we observed large variations in performance across sessions, the model based its choices on session-gain modulated evidence *<u>S</u>*_*ti*_ = *γ*_*t*_*S*_*ti*_ instead raw evidence *S*_*ti*_ (this had no impact for the deterministic variant since *<u>S</u>*_*ti*_ and *S*_*ti*_ always have the same sign). We fitted the model from individual subject responses either with lapses *π*_*L*_and *π*_*R*_ as free parameters, or fixed to *π*_*L*_ = *π*_*R*_ = 0.01. Figures in the main manuscript correspond to the deterministic snapshot model with fixed lapses. We also studied variants of the snapshot model where decisions in each trial are based on *K* attended samples, i.e depends on (*S*_*ti*_,.. *S*_*t,i*+*K*−1_) with 1 ≤ *K* ≤ *n* − 1 and 1 ≤ *i* ≤ *n* − *K* + 1 is the first attended sample. In the deterministic case, the choice is directly determined by the sign of the sum of the signed evidence for the attended samples. In the non-deterministic case, the evidence for the attended samples are weighted and passed through a sigmoid: *H*_*i*_(*S*_*t*_) = *σ*(*Σ*_*k*∈[1..*K*]_*β*_*ki*_*S*_*t,i*+*k*−1_). The model with a single attended sample presented above is equivalent to this extended model when using *K* = 1. At the other end, using *K* = *n* corresponds to the temporal integration model (without the lateral bias).

### Extrema-detection model

In the extrema-detection model, a choice is selected according to the first sample in the sequence whose absolute evidence value reaches a certain threshold *θ*, i.e *p*(*r*_*t*_|*θ*) = *H*(*m*_*ti*_), |*m*_*ti*_| ≥ *θ*, |*m*_*tj*_| < *θ* for all *j* < *i*. Here *m*_*ti*_ is the sample evidence corrupted by sensory noise *ε*_*ti*_ which is distributed normally with variance *σ*^2^: *m*_*ti*_ = *S*_*ti*_ + *ε*_*ti*_ with *ε*_*ti*_ ∼ *N*(0, *σ*^2^). *H* is the step function. If the stimulus sequence ends and no sample has reached the threshold, then the decision is taken at chance. As described in (Waskom and Kiani 2018), the probability for a rightward choice at trial *t* can be expressed as:

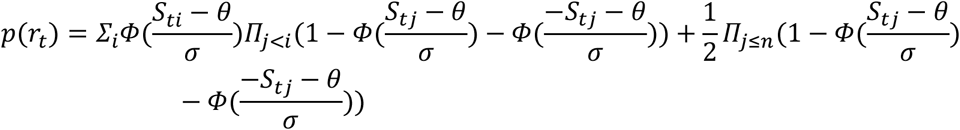

where *Φ* is the cumulative normal distribution. We also included the possibility for left and right lapses with probability *π*_*L*_and *π*_*R*_. Following Stine and colleagues (Stine et al. 2020), we explored an alternative default rule called *‘*last sample*’* rule: if the stimulus extinguishes and the threshold has not been reached, then the decision is based on the (noisy) last sample rather than simply by chance. This changes the equation describing the probability for rightward choices to:

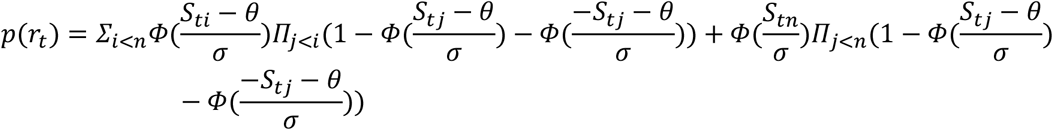

As for the snapshot model, we used the session-gain modulated evidence *<u>S</u>*_*ti*_ instead of raw evidence *S*_*ti*_ for fitting the model to monkey and rat data. The four parameters of the model (*θ, σ, π*_*L*_, *π*_*R*_) were estimated from each subject data by maximizing the log-likelihood with interior-point algorithm (function *fmincon* in Matlab) and 10 different initializations of the parameters.

### Model validation and model comparison

Psychophysical kernels were obtained from subject data and simulated data by running a logistic regression model: *p*(*r*_*t*_) = *σ*(*β*_0_ + *Σ*_*i*_*β*_*i*_*S*_*ti*_). Standard errors of the weights *β*_*i*_were obtained from the Laplace approximation. For psychometric curves, we first defined the weighted stimulus evidence *T*_*t*_ at trial *t* as the session-modulated weighted sum of signed sample evidence; with the weights obtained from the logistic regression model above *T*_*t*_ = *γ*_*t*_*Σ*_*i*_*β*_*i*_*S*_*ti*_. We then divided the total stimulus evidence into 50 quantiles (10 for human subjects) and computed the psychometric curve as the proportion of rightward choices for each quantile.

The boundary performance for the snapshot and extrema-detection models corresponds to the best choice accuracy out of all the parameterizations for each model. In the snapshot model, the boundary performance corresponds to the deterministic version with no-lapse, where the attended sample is always the sample *i*^*^whose sign better predicts the stimulus category over all animal trials, i.e. *π*_*i*_* = 1 and *π*_*i*_ = 0 if *i* ≠ *i*^*^. For the extrema-detection model, the boundary performance corresponds to the lapse-free model with no sensory noise (*σ* = 0) and a certain value for threshold *θ* that is identified for each subject by simple parameter search.

Finally, model selection was performed using the Akaike Information Criterion *AIC* = 2*p* − 2*L*_*ML*_, where *p* is the number of model parameters and *L*_*ML*_ is the likelihood evaluated at maximum likelihood parameters.

### Analysis of majority-driven choices

We selected for each animal the subset of trials corresponding to when the largest evidence sample was at odds with the total stimulus evidence, i.e. where *sign*(*S*_*tj*_, |*S*_*tj*_| ≥ |*S*_*ti*_| v *i*) ≠ *sign*(*Σ*_*i*_*S*_*ti*_). For this subset of trials, we computed the proportion of animal choices that were aligned with the overall stimulus evidence. We repeated the analysis for simulated data from the integration and extrema-detection models.

### Subjective weighting analysis

In order to estimate the impact of each sample on the animal choice as a function of sample evidence, we built and estimated the following statistical model

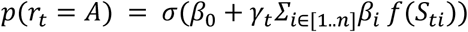

As can be seen, this model is equivalent to the temporal integration model under the assumption that *f* is a linear function. Rather, here we wanted to estimate the function *f* (as well as the session gain *γ*_*t*_, lateral bias *β*_0_and sensory weight *β*_*i*_). Including the session gain was necessary for estimating *f* accurately from the monkey and rat behavioral data, since the distribution of pulse strength *S*_*ti*_ was varied across sessions and could otherwise induce a confound. We assumed that *f* is an odd function, i.e. *f*(−*S*_*ti*_) = −*f*(*S*_*ti*_). This equation takes the form of a Generalized Unrestricted Model and was fitted using the Laplace approximation method as described in (Adam and Hyafil 2020). In the monkey experiment, sample evidence could take only a finite number of values, so *f* was simply estimated over these values. In the human experiment, sample evidence could take continuous values. In this case, we defined a Gaussian Process prior over *f* with squared exponential kernel with length scale 0.1 and variance 1.

### Integration of early and late evidence

We designed a new analysis tool to characterize the statistical mapping from the multidimensional stimulus space ***S***_*t*_ = (*S*_*t*1_, … *S*_*tn*_) ∈ ℜ^*n*^onto binary choices *r*_*t*_ ∈ [0,1]. We first collapsed the stimulus sequence ***S***_*t*_ onto the two-dimensional space defined by early evidence ***E***_*t*_ and late evidence ***L***_*t*_ defined by *E*_*t*_ = *γ*_*t*_*Σ*_1≤*i*≤[*n*/2]_*β*_*i*_*S*_*ti*_ and *L*_*t*_ = *γ*_*t*_*Σ*_[*n*/2]+1≤*i*≤*n*_*β*_*i*_*S*_*ti*_, where the weights *β*_*i*_ and session gains *γ*_*t*_correspond to parameters estimated from the temporal integration model (session gains were omitted for human participants). Next we plotted the integration map which represents the probability for rightward choices as a function of (*E*_*t*_, *L*_*t*_). The map was obtained by smoothing data points with a two-dimensional gaussian kernel. More specifically, for each pair value *(E,L)*, we selected the trials whose early and late evidence values *E*_*t*_ and *L*_*t*_ fell within a certain distance to *(E,L)*, i.e. *d*_*t*_ = *dist*((*E, L*)(*E*_*t*_, *L*_*t*_)) < We then computed the proportion of rightward choices for the selected trials, with a weight for each trial depending on the distance to the pair value *w*_*t*_ = *N*((*E*_*t*_, *L*_*t*_); (*E, L*),0. 1^2^ *I*). Because the space *(E,L)* was not sampled uniformly during the experiment, we represent the density of trials by brightness. For each subject we obtained integration maps both from subject data as well as from model simulations. For each model, we computed the Pearson correlation between the maps obtained from the corresponding simulation and from the subject data. We tested the significance of correlation measures between models by using a bootstrapping procedure: we calculated the correlation measure *r* from 100 bootstraps for each model and participant, and then performed an unpaired t-test between bootstrapped *r*.

Next, we analyzed the conditional psychometric curves, i.e. the psychometric curves for the early evidence conditioned on the value of late evidence, which correspond to vertical cuts in the integration map. To do so, we first binned late evidence *L*_*t*_ by bins of width 0.5. Conditional psychometric curve represent the probability of rightward choices as a function of early evidence *E*_*t*_, separately for each late evidence bin. For each late evidence bin, we also estimated the corresponding bias *β*, left lapse *π*_*L*_ and right lapse *π*_*R*_ by fitting the following function on the corresponding subset of trials:

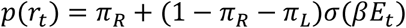

### Analysis of LIP neuron activity

We analyzed the activity of 82 LIP neurons recorded over 43 sessions of the motion discrimination tasks (Yates et al. 2017). We applied the following procedure to extract the integration map for LIP neurons. For each neuron *n*, we computed the spike count *s*_*t*_^(*n*)^ in a window of 500 ms width following each stimulus offset, which is where LIP neurons were found to have maximal selectivity to motion evidence from the entire pulse sequence (Yates et al. 2017). We then applied a Poisson GLM *E*(*s*_*t*_^(*n*)^) = *exp*(*w*_0_^(*n*)^ + *Σ*_*i*_*w*_*i*_^(*n*)^*S*_*ti*_) for each neuron *n* to extract the impact of each sample *i* on the individual neural spike count *w*_*i*_^(*n*)^. For each trial *t*, we used these weights to compute the neuron-weighted early and late evidence defined by and *E*_*t*_^(*n*)^ = *Σ*_1≤*i*≤3_*w*_*i*_^(*n*)^*S*_*ti*_ *L*_*t*_^(*n*)^= *Σ*_4≤*i*≤7_*w*_*i*_^(*n*)^*S*_*ti*_. Note that this weighting converts the evidence onto the space defined by the preferred direction of the neuron, such that positive evidence signals evidence towards the preferred direction and negative evidence signals evidence towards the anti-preferred direction. We then merged the vectors for normalized spike counts *<u>s</u>*_*t*_^(*n*)^ = *s*_*t*_^(*n*)^/*exp*(*w*_0_^(*n*)^), early evidence *E*_*t*_^(*n*)^ and late evidence *L*_*t*_^(*n*)^ across all neurons. The normalized spike counts were binned by values of early and late evidence (bin width: 0.02), and the average over each bin was computed after convolving with a two-dimensional gaussian kernel of width 0.1. The neural integration map represents the average normalized activity per bin.

Simulations of spiking data for the integration and non-integration models were proceeded as follows. First, the neural integration model corresponds to linear summing with neuron-specific weights which are then passed through an exponential nonlinearity; the spike counts for each trial are generated using a Poisson distribution whose rate is equal to the nonlinear output (Supp Figure 9a, top). This corresponds exactly to the generative process of the Poisson GLM described above. For the extrema detection model (Supp Figure 9a middle), we hypothesized that LIP activity would only be driven by the sample that reaches the threshold (and dictates the animal response). To this end, we first simulated the behavioral extrema detection model for all trials, using parameters (*θ, σ, π*_*L*_, *π*_*R*_) fitted from the corresponding animal, to identify which sample *i* reaches the subject-specific threshold. We then assumed that the spiking activity of the neuron would follow the stimulus value at sample *i S*_*ti*_ (signed by the preferred direction of the neuron *p*^(*n*)^ through:

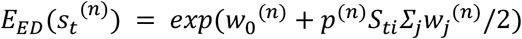

Again the spike count were generated from a Poisson distribution with rate *E*_*ED*_(*s*_*t*_^(*n*)^).

Finally, for the snapshot model (Supp. Figure 9a bottom), we assumed that the neuron activity would merely reflect the sensory value of the only sample it would attend. We assumed that the probability mass function to attend each of the 7 samples would be neuron-specific, so we used the normalized weights of the Poisson GLM for that specific neuron as defining such probability (weights were signed by the neuron preferred direction so that the vast majority of weights were positive; negative weights were ignored). For each trial, we thus randomly sampled the attended sample *i* using this probability mass function and then simulated the spike count *s*_*t*_^(*n*)^ from a Poisson distribution with rate *E*_*Snapshot*_(*s*_*t*_^(*n*)^) = *exp*(*w*_0_^(*n*)^ + *p*^(*n*)^*S*_*ti*_*Σ*_*j*_*w*_*j*_^(*n*)^).

We simulated spiking activity for each neuron and for each integration and non-integration model, and then used simulated data to compute neural integration maps exactly as described above for the actual LIP neuron activity.

## Data and code availability

All experimental data (behavioral and neural data in monkeys, behavioral data in rats and humans) and code to run the analysis will be made publicly available at https://github.com/ahyafil prior to final publication

## ACKNOWLEDGMENTS

The authors thank Jake Yates for sharing information regarding the monkey experimental data. The authors are supported by the Spanish State Research Agency (RYC-2017-23231 to A.H.), Spanish Ministry of Economy and Competitiveness together with the European Regional Development Fund grant SAF2015-70324-R (to JR), European Research Council grant ERC-2015-CoG-683209 (to JR), NIH grant R01EY017366 (to ACH & JWP), NIH BRAIN Initiative grant NS104899 (to JWP) and the Simons Collaboration on the Global Brain (SCGB AWD543027, JWP).

## SUPPLEMENTARY FIGURES

**Supplementary Figure 1.**
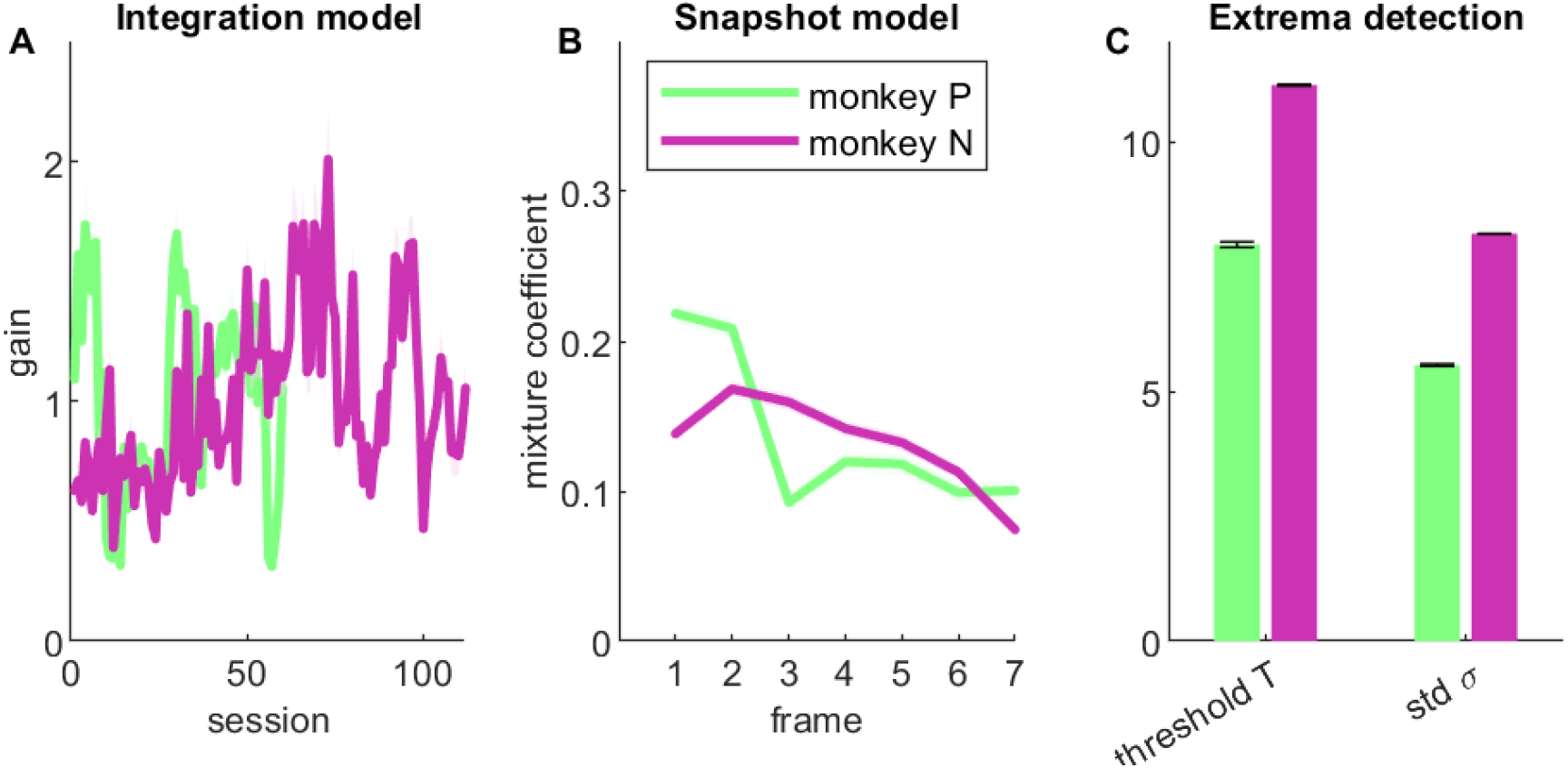
Parameter fits for integration and non-integration models. **A**. Modulation gain *γ* per session for the integration model, for each animal (green: monkey P; purple: monkey N). **B**. Mixture coefficients *π*_*i*_ of the snapshot model estimated for each monkey, representing the prior probability that each sample is attended on each trial. **C**. Parameters *T* and *σ* of the extrema-detection model, estimated for each monkey. Error bars correspond to the confidence interval obtained using the Laplace approximation.

**Supplementary Figure 2.**
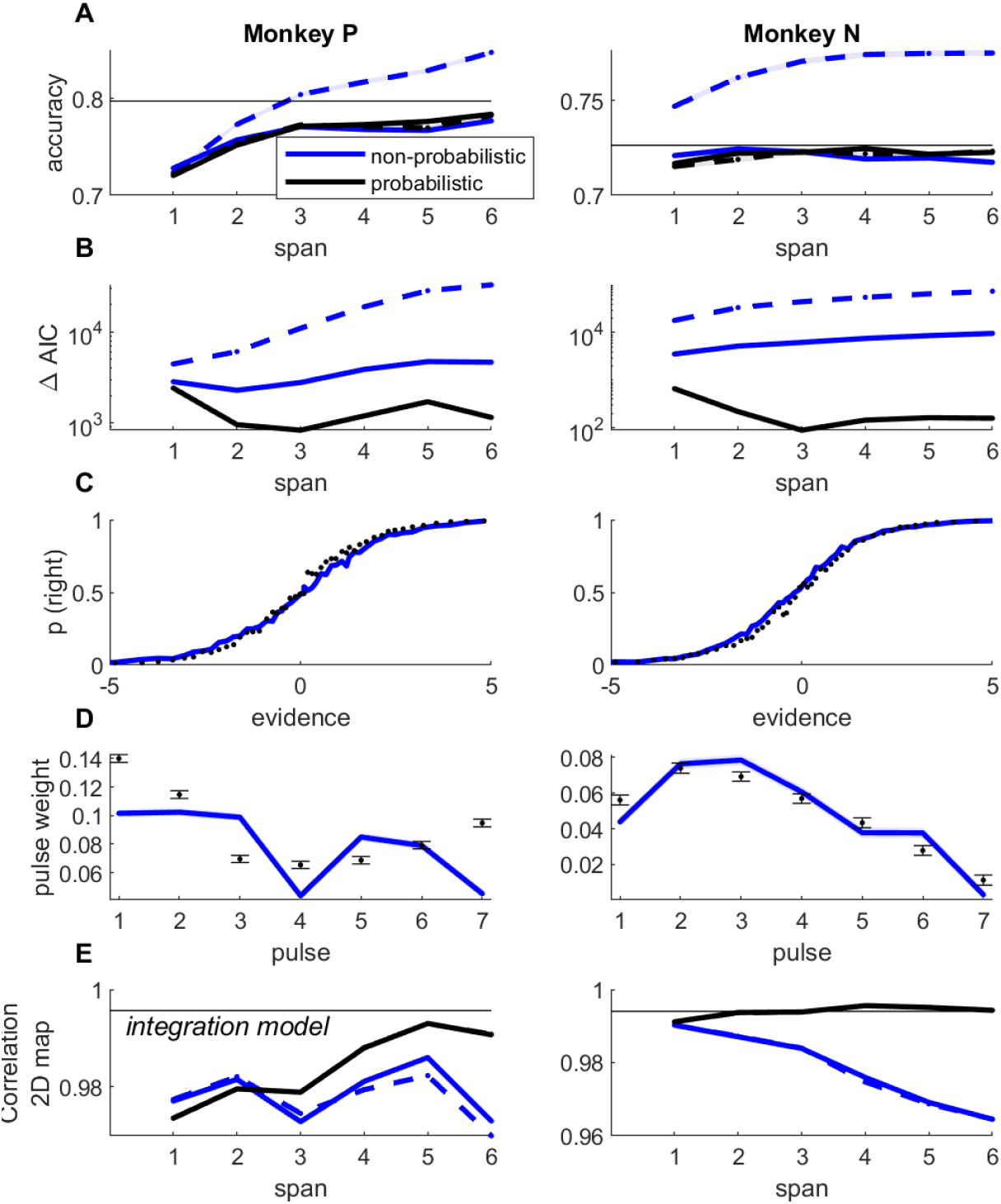
Model fits for variants of the snapshot model. **A**. Predicted accuracy for the snapshot model fitted to monkey data, as a function of memory span *K*, for fixed lapses (blue curve, *π*_*L*_ = *π*_*R*_ = 0.01) and lapses estimated from the data (black curve). Full lines represent the model with sensory noise (*“*probabilistic*”*), dotted lines represent the model without sensory noise (*“*non-probabilistic*”*). Memory span *K* corresponds to the number of successive samples used to define the decision on each trial (see Methods). The horizontal bar corresponds to the average accuracy of the animal. **B**. AIC difference between snapshot and integration model. Legend as in A. Positive values indicate that the snapshot model provides a worse fit. **C**. Psychometric curve for the snapshot model with span *K=3* samples, sensory noise and free lapse parameters (best snapshot model variant according to AIC). **D**. Psychophysical kernel for the same variant of the model. **E**. Correlation between data and model integration maps for variants of the snapshot model.

**Supplementary Figure 3.**
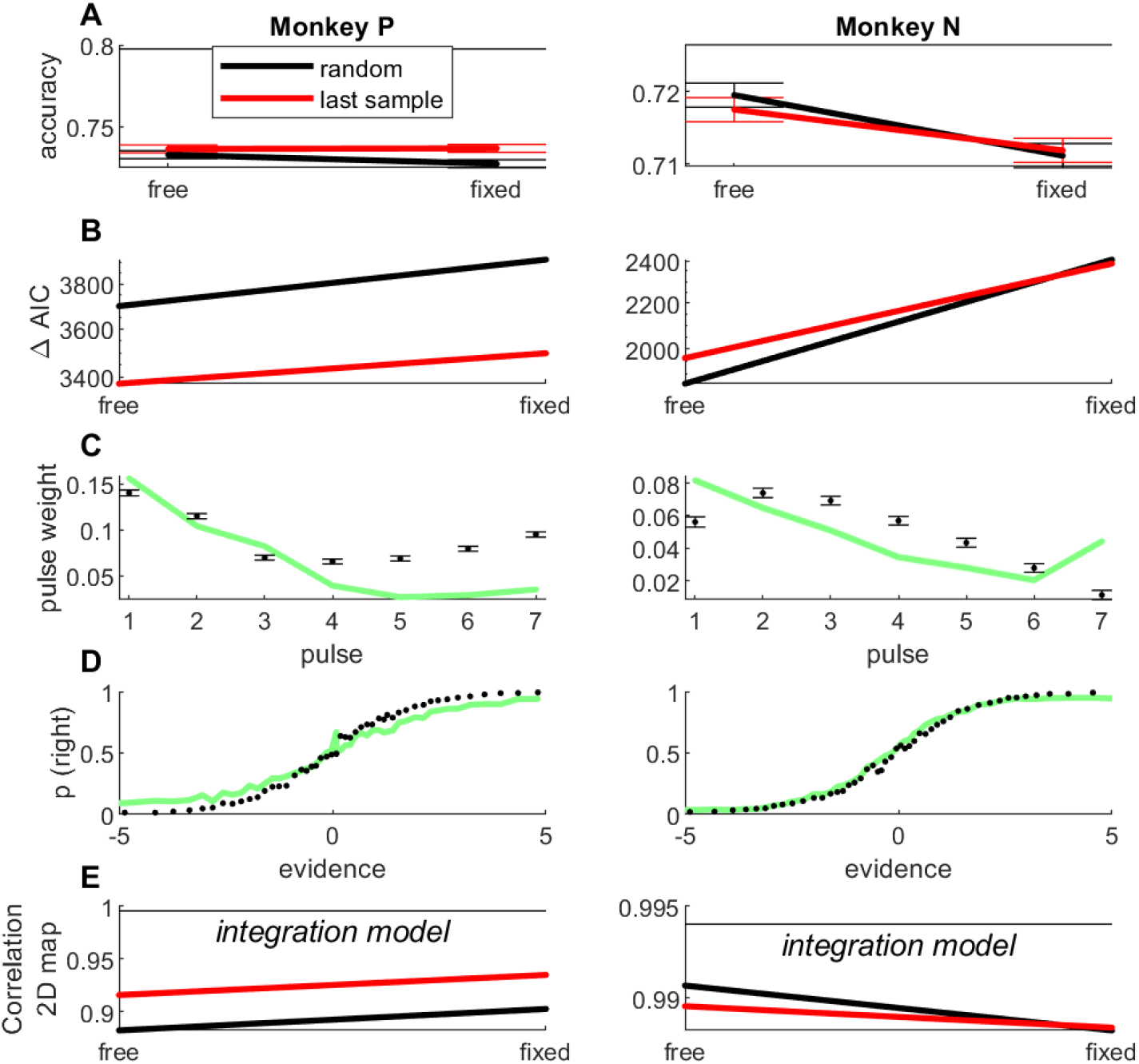
Model fits for variants of the extrema-detection model. **A**. Predicted accuracy for the extrema-detection model fitted to the monkey data, for random (black curves) and last sample (red curve) default rule, and for fixed lapses (*π*_*L*_ = *π*_*R*_ = 0.01) or lapse parameters estimated from the data. The horizontal bar indicates animal accuracy. **B**. AIC difference between variants of the extrema-detection model and the integration model. Legend as in A. Positive values indicate that the extrema-detection model provides a worse fit. **C-D**. Psychometric curve (C) and psychophysical kernel (D) for the model variant that provided the best match to behavior in terms of predicted accuracy and AIC: free lapse parameters and last sample rule. **E**. Correlation between integration maps from animal data and simulated data (see Figure 4) for variants of the extrema-detection model. The horizontal bar marks the correlation between experimental data and the integration model.

**Supplementary Figure 4.**
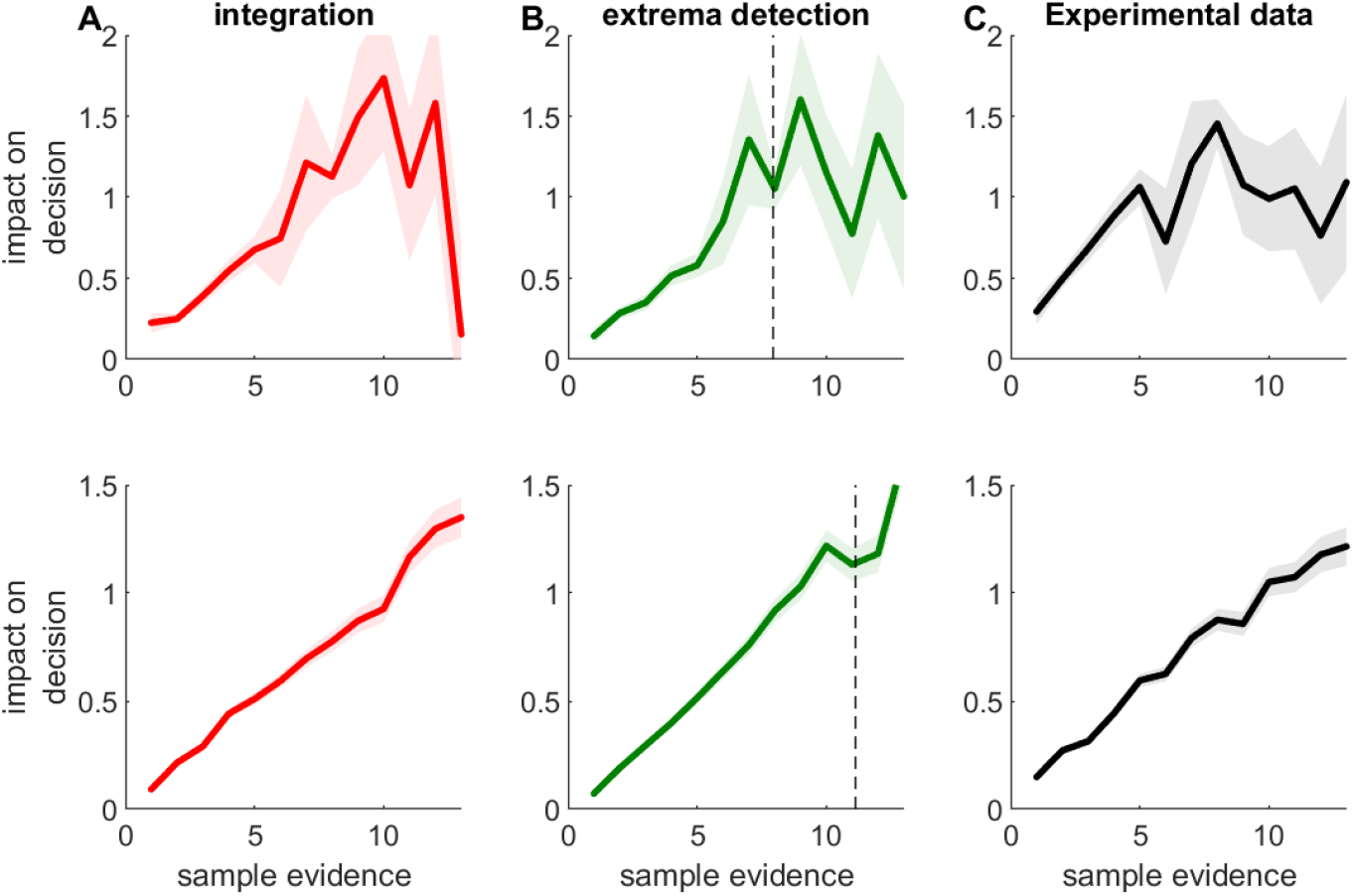
Subjective weights for animal data and simulated models. Impact on decision of individual samples as a function of absolute sample evidence. Shaded area: standard error of the weight. Top row: monkey P; bottom row: monkey N. **A**. Integration model. **B**. extrema-detection model. The vertical dotted line marks the value of the threshold *T* estimated from animal data. **C**. Impact on decision of individual pulses, estimated from each monkey.

**Supplementary Figure 5.**
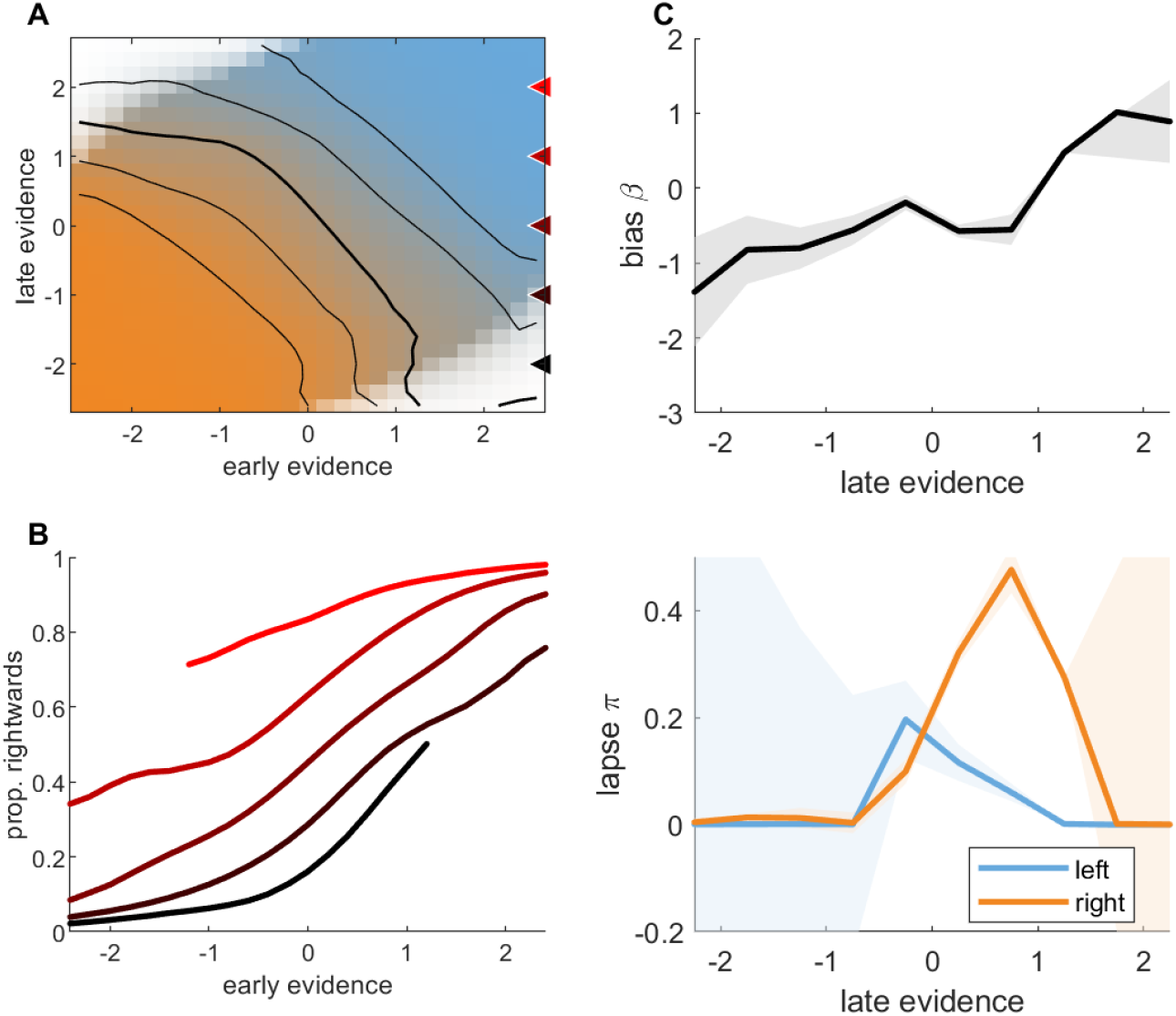
Integration of early and late evidence for monkey P. **A**. Integration map. Legend as in Figure 4A. **B**. Conditional psychometric curves. Legend as in Figure 4B. **C**. Bias and lapse parameters from conditional psychometric curves, as a function of late evidence. Legend as in Figure 4D-E.

**Supplementary Figure 6.**
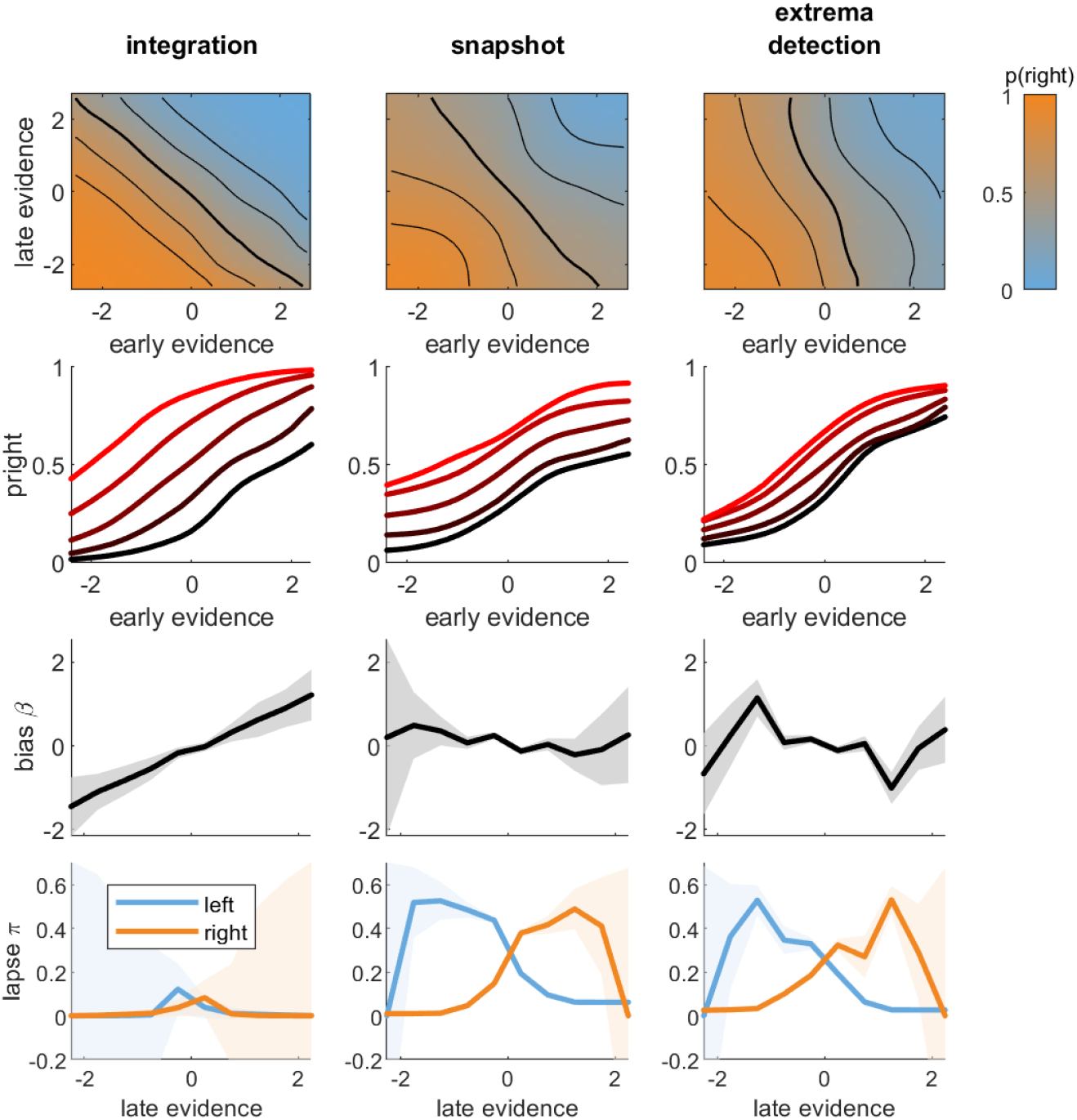
Integration between early and late evidence for simulated data from integration and non-integration models. Data was simulated for each model from parameters estimated from monkey N. Left panels: integration model. Middle panels: snapshot models. Right panels: extrema-detection models. **A**. Integration maps. **B**. Conditional psychometric curves. **C**. Lateral bias and **D**. lapse parameters estimated from conditional psychometric curves, as a function late evidence. Legend as in Figure 4.

**Supplementary Figure 7.**
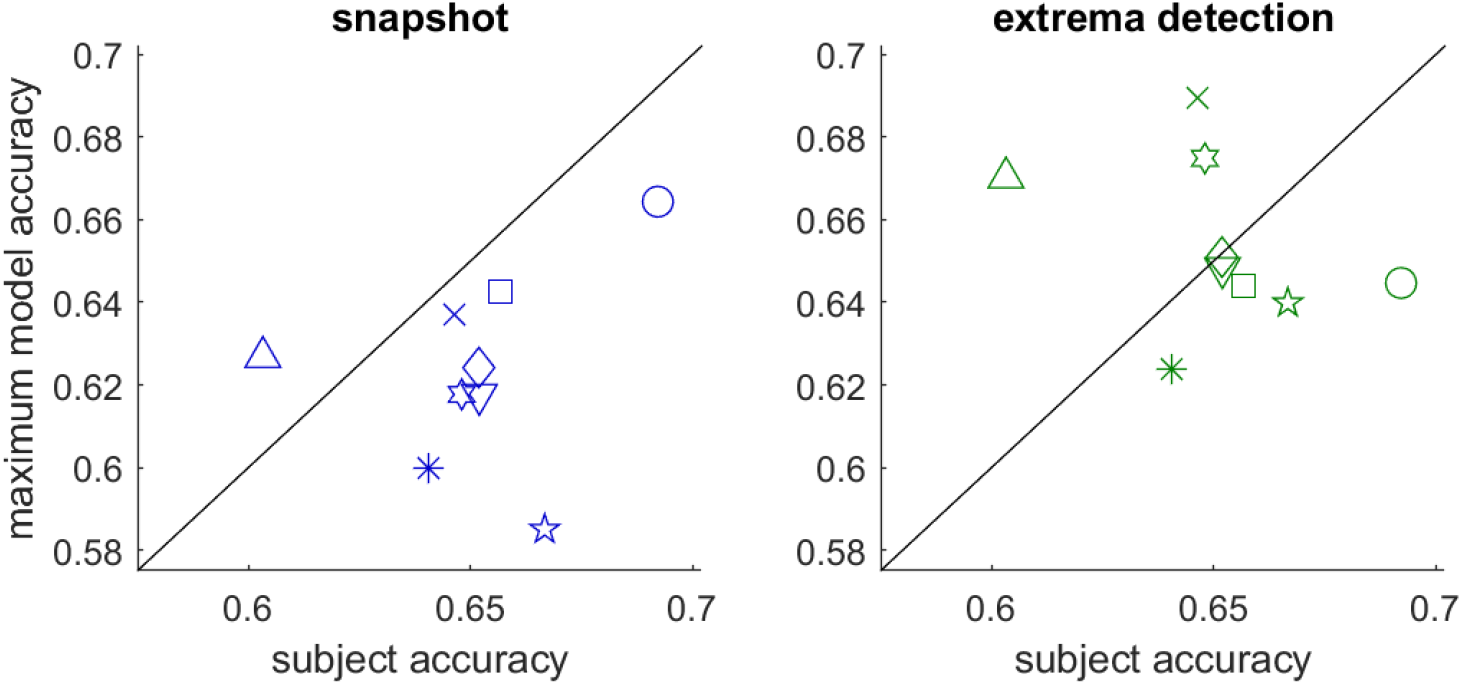
Maximum accuracy of the non-integration models vs. human subject accuracy in the orientation discrimination task. Left panel: snapshot model (with span *K*=1). Right panel: extrema-detection. Each symbol represents a subject.

**Supplementary Figure 8.**
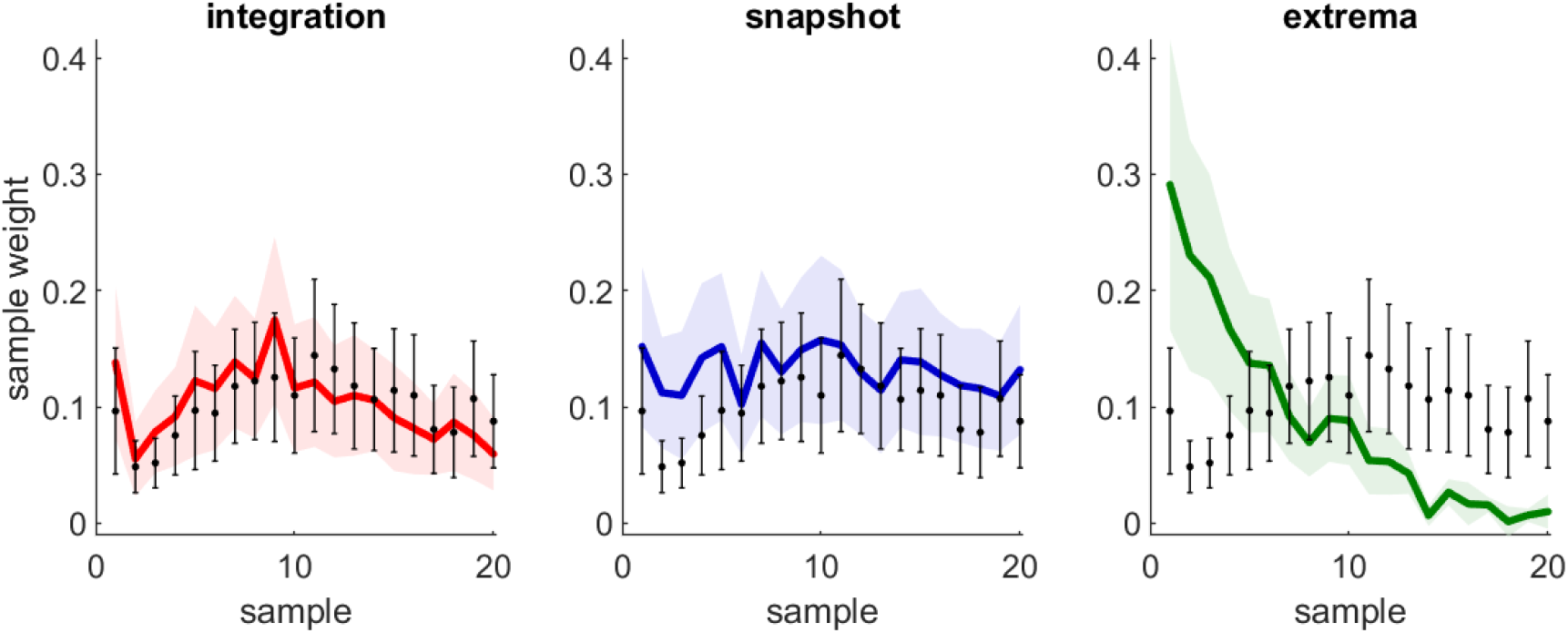
Psychophysical kernels for animals and models in rats (*n*=3) performing the DSS task with 20-sample stimuli.

**Supplementary Figure 9.**
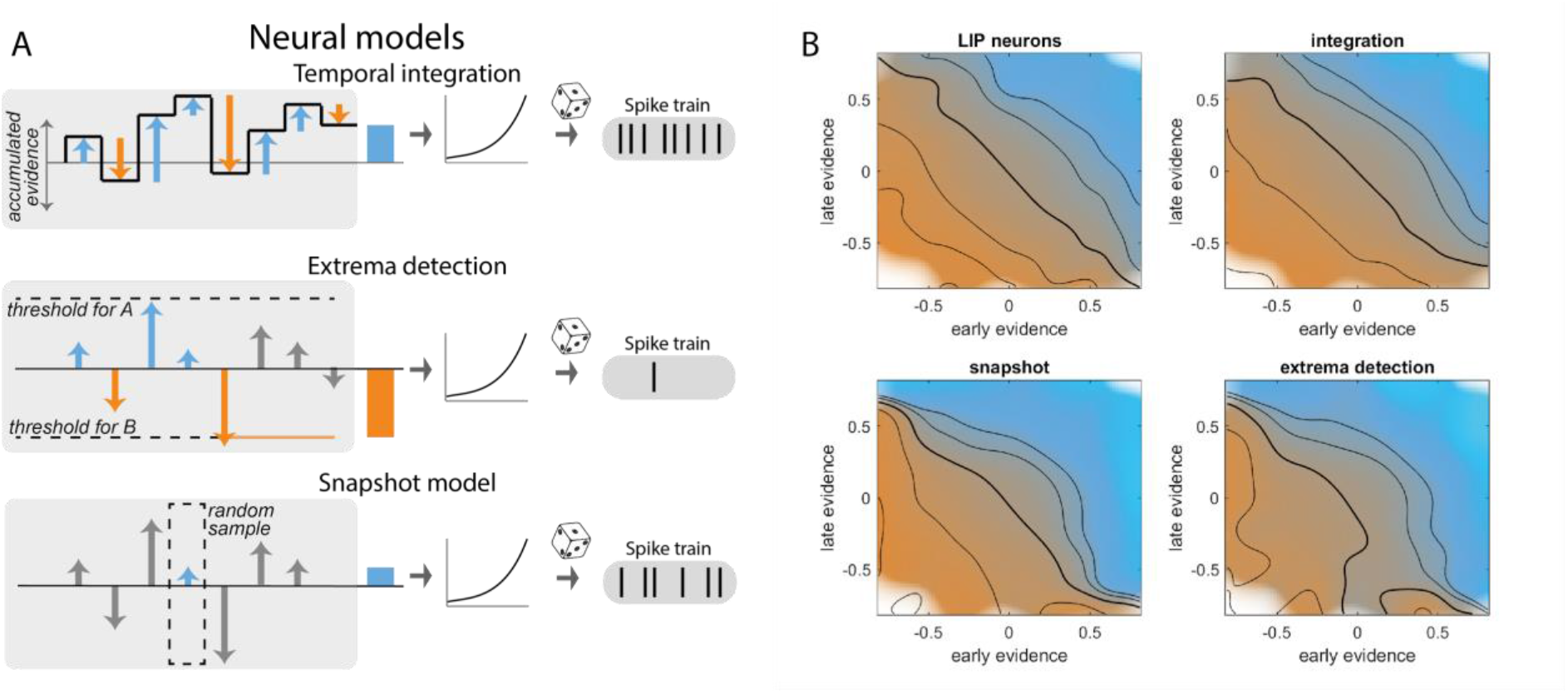
Individual LIP neurons integrate sensory information over stimulus sequence. **A**. Neural models for temporal integration, extrema-detection and snapshot model. **B**. Integration map for LIP neurons, and simulated neurons following either integration, extrema-detection or snapshot model. Color represents the average normalized spike count per bins of neuron-weighted early and late evidence (see Methods). Isolines represent values of 0.4, 0.6, 1, 1.4 and 1.8.

